# Auranofin induces lethality driven by reactive oxygen species in high-grade serous ovarian cancer cells

**DOI:** 10.1101/2023.09.13.557629

**Authors:** Farah H. Abdalbari, Elvis Martinez-Jaramillo, Benjamin N. Forgie, Estelle Tran, Edith Zorytcha, Alicia A. Goyeneche, Carlos M. Telleria

**Affiliations:** Experimental Pathology Unit, Department of Pathology, Faculty of Medicine and Health Sciences, McGill University, Montreal, QC H3A 2B4, Canada; Cancer Research Program, Research Institute, McGill University Health Centre, Montreal, QC H4A 3J1, Canada

**Keywords:** Auranofin, high grade serous ovarian cancer, TrxR, apoptosis, DNA damage, ROS, L-buthionine sulfoximine, cisplatin, N-acetyl cysteine, drug repurposing, GSH

## Abstract

High-grade serous ovarian cancer (HGSOC) accounts for 70% of ovarian cancer cases in the clinical setting; the survival rate for this disease remains remarkably low due to the lack of long-term consolidation therapies following the standard platinum-based chemotherapy, which is not long lasting as the disease recurs as platinum resistant. The purpose of this study was to explore a novel treatment against HGSOC using the gold complex auranofin (AF). AF primarily functions as a pro-oxidant agent through the inhibition of thioredoxin reductase (TrxR), an antioxidant enzyme that is overexpressed in ovarian cancer. We investigated the effect of AF on TrxR activity and various mechanisms of cytotoxicity using HGSOC cells that are clinically sensitive or resistant to platinum. In addition, we studied the interaction between AF and another pro-oxidant agent, L-buthionine sulfoximine (L-BSO), an anti-glutathione (GSH) compound. We demonstrate that AF potently inhibits TrxR activity and reduces the vitality and viability of HGSOC cells regardless of their sensitivities to platinum. We show that AF induces accumulation of reactive oxygen species (ROS), triggers the depolarization of the mitochondrial membrane, and kills HGSOC cells by inducing apoptosis. Yet, AF-induced cell death is abrogated by the ROS-scavenger N-acetyl cysteine (NAC). In addition, the lethality of AF is associated with the activation of caspases-3/7 and the generation of DNA damage, effects that are also prevented by the presence of NAC. Finally, when AF and L-BSO are combined, we observed a synergistic lethality against HGSOC cells, which is mediated by a further increase in ROS together with a decrease in the levels of the antioxidant GSH. In summary, our results support the concept that AF can be used alone or in combination with L-BSO to kill HGSOC cells regardless of their platinum sensitivities suggesting that depletion of antioxidants is an efficient route for treating this disease.

## Introduction

Ovarian cancer remains the eight-leading cause of cancer-related deaths among women worldwide [1]. According to GLOBOCAN, there were 313,959 new ovarian cancer cases and 207,252 deaths due to ovarian cancer in 2020 [2]. Over the past few decades, there has been a small reduction in the incidence rates of ovarian cancer, primarily due to preventative measures, such as the introduction of oral contraceptives [3], and the decline in the use of menopausal hormonal therapy [4]. However, there has been minimal improvement in the overall survival of patients carrying the disease [1,5]. Treatment with platinating agents is very efficient at the beginning of the illness following diagnosis and debulking surgery, leading to an initial response of 80% [6,7]. However, over time the disease almost always recurs with a platinum-resistant phenotype that is highly difficult to treat, thus explaining its high mortality [8]. Acquired platinum resistance (either to cisplatin or carboplatin) occurs via multipronged mechanisms, including the decreased intracellular accumulation of the compounds, increased drug detoxification, and increased activity of the DNA repair machinery among the most often investigated [5].

To counteract the high mortality of ovarian cancer it is evident that platinum-based chemotherapy needs to be coupled with a follow up therapy that should be used chronically to maintain the disease in a dormant stage. We propose that one such approach is the chronic use of drugs with cytotoxic activity that could be repurposed from other diseases. Drug repurposing is a cost-effective approach in which drugs that have been approved for one use are repositioned for a different use than the one they were originally approved for [9,10]. For instance, our laboratory has shown that the antiprogesterone/anti-glucocorticoid agent mifepristone is efficient as upfront therapy or after cisplatin and/or paclitaxel therapy against ovarian cancer cells [5,11-14]. We have also shown the efficacy of the HIV inhibitor nelfinavir against ovarian cancer cells sensitive or resistant to platinum [15]. Here, we are adding auranofin, a gold complex approved in 1985 to treat rheumatoid arthritis [16] to the list of anti-ovarian cancer drugs emerging from repurposing auranofin.

The primary mechanism of action of auranofin is to act as a pro-oxidative agent, increasing the production of reactive oxygen species (ROS) as a consequence of inhibiting the thioredoxin reductase (TrxR) anti-oxidant system [17]. TrxR has been found overexpressed in various cancers, including non-small cell lung cancer [18], breast cancer [19], and cisplatin-resistant ovarian cancer [20]. The TrxR system is involved in the overall promotion of tumor progression by preventing cell death triggered by oxidative stress [21]. Of interest, TrxR overexpression is associated with short overall survival in ovarian cancer patients based on a Kaplan Meir survival analysis [21]. These findings suggest that TrxR is an attractive therapeutic target against ovarian cancer, and auranofin a potent TrxR inhibitor and pro-oxidative agent that can be used to combat this disease. Previous reports on various cancer cells have demonstrated that auranofin induces inhibition of cell proliferation by causing overproduction of ROS [22], caspase-independent apoptosis [23], and cell death triggered by DNA damage [24]. Additionally, auranofin has been shown to inhibit angiogenesis [25], protein homeostasis [26,27], and deubiquitinases involved in proteasome-mediated protein degradation [28]. These findings signify that auranofin is a potent anti-cancer agent that negatively targets multiple metabolic pathways of the cancer cell. In this study, we identify the mechanisms of cytotoxicity induced by auranofin in HGSOC cells that have different clinical sensitivities to platinum. We show that auranofin causes ROS-dependent inhibition of cell proliferation, caspase-associated apoptosis, mitochondrial membrane depolarization, DNA damage, cleavage of poly-ADP ribose polymerase (PARP), and the poly-ubiquitination of proteins [28]. Additionally, we show a synergistic lethal interaction between auranofin and a second pro-oxidant agent, the glutathione (GSH) inhibitor, L-buthionine sulfoximine (L-BSO); this drug interaction involving two blockers of key antioxidant pathways that cancer cells rely upon is dependent on the presence of ROS.

## Materials and Methods

### Reagents and Cell Lines

PEO1 cells are epithelial ovarian cancer cells isolated from a patient after its first relapse 22 months following treatment with cisplatin, 5-fluorouracil, and chlorambucil, while the patient was still sensitive to platinum-based chemotherapy; PEO4 cells were subsequently isolated from the same patient after the second relapse in which the patient was no longer sensitive to the chemotherapy. These cell lines were histologically characterized in 1988 and sequenced in 2010 [29,30], whereas we authenticated them in 2020 based on their autosomal short-tandem repeats [15]. The cells were cultured in RPMI 1640 media (Mediatech, Manassas, VA, USA) supplemented with 5% fetal bovine serum (FBS) (Atlanta Biologicals, Lawrenceville, GA, USA), 5% bovine serum (Life Technologies, Auckland, New Zealand), 0.01 mg/mL of human insulin (Roche, Indianapolis, IN, USA), 10 mM HEPES (Corning, Corning, NY, USA), 100 IU penicillin (Mediatech), 100 μg/mL streptomycin (Mediatech), 2 mM L-Alanyl-L-Glutamine (Glutagro^TM^, Corning) and 1 mM sodium pyruvate (Corning). The cells were incubated in a 37 °C humidified incubator with 5% CO_2_. The drugs used in this study include: auranofin (Sigma Chemical Co., St. Louis, MO, USA), bortezomib (BZ) (Velcade®, Millennium Pharmaceuticals, Cambridge, MA, USA), L-buthionine sulfoximine (L-BSO) (Sigma), and N-acetyl cysteine (NAC; Sigma).

### Cellular Vitality

To determine the wellbeing of the cells, we performed a cellular vitality assay [31] (not viability) as we did not assess life versus dead cells but instead the health of the mitochondrial function by evaluating their enzyme activities as a surrogate marker of drug toxicity. PEO1 cells and PEO4 cells growing at 70% confluency were harvested and seeded in triplicate in 96-well plates at a density of 2.5x10^3^ cells/well in HGSOC medium and allowed to adhere overnight at 37°C in 5% CO_2_. The cells were then treated with auranofin at varying concentrations for 72 hours. Cell vitality was measured by adding 10 μl/well of 5 mg/ml MTT [3-(4, 5- dimethylthiazol-2-yl)-2, 5-diphenyltetrazolium bromide] (Life Technologies, Burlington, ON, Canada) in PBS solution. Cells were incubated for 4 hours at 37°C in 5% CO_2_ where the tetrazolium dye is reduced to insoluble formazan. One hundred microliters/well of 10% sodium dodecyl sulfate (SDS)/ 0.01 M HCl was added to stop the reaction. The absorbance was recorded at 570 nm following overnight incubation. Blank controls were subtracted and % cell vitality relative to the control was calculated.

### Cellular Viability

Clinically platinum-sensitive (PEO1) and clinically platinum-resistant (PEO4) HGSOC cells were treated with increasing concentrations of auranofin for 72 hours. The cells were collected, centrifuged, and the remaining cell pellet was resuspended in 1 mL of cell culture medium. An aliquot of cells was taken and stained with the Muse® count and viability reagent (Luminex, Austin, TX, USA) for 5 minutes; this reagent contains a DNA-binding dye that tags nucleated cells, and a second dye that differentiates live from dead cells by penetrating cells with compromised membrane integrity (i.e., non-viable cells). Stained cells were analyzed using the Muse® micro-capillary cytometer (Millipore, Hayward, ON, Canada), and viable cell number and total cell number were determined.

### Clonogenic Survival

To assess the residual toxicity of auranofin on HGSOC cells, 1000 viable cells were taken from the cell culture treated with auranofin for 72 hours and plated in media devoid of drug in a 6-well plate for a period of 2 weeks. The experiment was terminated once the vehicle-treated group contained positive colonies. Positive colonies refer to colonies that contain 50 or more cells; this is used as a measure of assessing the proliferative capacity of the cells in a long-term period even though they survived the initial 72 hours of drug treatment; in other words, we tested how the exposure to auranofin affected their long-term reproductive capacity.

### Cell Cycle Distribution

Following the 72-hour treatment with auranofin, PEO1 cells and PEO4 cells were fixed using 4% paraformaldehyde (PFA) and stored at 4° C overnight. Samples were centrifuged, and the cell pellet was washed with 500 µL of 1 X phosphate-buffered saline (PBS) (Corning, Manassas, VA, USA). Two hundred thousand cells were taken and centrifuged at 2000 x *g* for 5 minutes. The supernatant was discarded, and the pellet was resuspended in 200 μL of cell cycle buffer containing 0.5 mg/mL propidium iodide, a cell permeable DNA intercalating agent that serves to analyze the status of DNA content. Cell cycle analysis was completed using the Muse® micro-capillary cytometer (Millipore). This method was described previously in detail [15].

### Protein Lysate Preparation and Western Blot Analysis

PEO1 cells and PEO4 cells were treated with increasing concentrations of auranofin for 72 hours, and whole cellular extracts were collected at the end of the incubation. The cells were centrifuged at 1500 x *g* for 6 minutes, the supernatant was removed, and the cell pellet was resuspended in 1 mL of cold 1 X PBS. The samples were centrifuged at 2,000 x *g* for additional 5 minutes, the supernatant was removed, and the cell pellets were snap frozen in liquid nitrogen and stored at -80°C until further processing. The proteins were isolated by first adding lysis buffer to the cell pellets. The lysis buffer was prepared by adding the following: 0.5% NP-40, 1 mM dithiothreitol (DTT), 1 mM phenylmethylsulphonyl fluoride (PMSF), 2 µg/mL aprotinin, 2 µg/mL pepstatin, 2 µg/mL leupeptin, 50 mM sodium fluoride, and 1 mM sodium orthovanadate. The cell pellets were resuspended in the lysis buffer by gentle vortexing, and were placed in ice on a shaker for 30 minutes at 4 °C. Samples were centrifuged at 12,000 x *g* for 15 minutes at 4°C. The proteins in the supernatant were collected and transferred to a separate tube. Protein samples were then quantified using the Pierce BCA Protein colorimetric assay purchased from Thermo Fisher Scientific (Rockford, IL, USA), and absorbance was measured at 562 nanometers using the Bio-Tek Cytation 3 Multi-Mode Reader (Agilent, Santa Clara, CA, USA). The proteins were run on a 10% SDS-polyacrylamide gel. Following the transfer onto the PVDF membranes, the membranes were blocked in 5% non-fat dry milk at room temperature for 1 hour and incubated with the primary antibodies at 4°C overnight. The membranes were washed 5 times in 1X TBS-T for 5 minutes each and incubated with the secondary antibody for 1 hour. The secondary antibody was removed, and the membranes were washed again 5 times in 1X TBS-T for 5 minutes each. The membranes were then imaged using the Bio-Rad ChemiDoc Imaging System (Bio-Rad, Hercules, CA, USA). Primary antibodies used were monoclonal anti-β-actin produced in mouse as clone AC-15 (A5442, Sigma), polyclonal anti-PARP produced in rabbit (9541, Cell Signalling Technology, Danvers, MA, USA), and polyclonal anti-ubiquitin produced in rabbit (3933S, Cell Signalling). Secondary antibodies were goat anti-rabbit IgG (H+L) conjugate (1706515, BioRad) and goat anti-mouse IgG (H+L)-HRP conjugate (1706516, BioRad).

### Detection of DNA Damage

To determine whether auranofin induces DNA damage in HGSOC, PEO1 cells and PEO4 cells were treated with 1, 2, or 4 µM of auranofin for 72 hours. Cells were collected and centrifuged at 300 x *g* for 5 minutes, and the supernatant was removed. The cells were resuspended in 50 µL of 1 X assay buffer per 100,000 cells, and equal volume of fixation buffer was added to the cells. The samples were incubated on ice for 10 minutes, spun down at 300 x *g* for 5 minutes, and the supernatant was discarded. The cells were resuspended in 90 µL of 1X assay buffer for every 100,000 cells. Cells were then stained with 10 µL of antibody working solution that was prepared by combining 5 µL of anti-phospho-ATM (Ser1981) labelled with phycoerythrin (PE), and 5 µL of anti-phospho-Histone H2A.X (Ser139) labelled with PE- Cyanine®5 (PeCy5). Samples were incubated at room temperature for 30 minutes protected from light. One hundred microliters of 1X assay buffer were added, and samples were centrifuged at 300 x *g* for 5 minutes. The supernatant was discarded, and the cells were resuspended in 200 µL of 1X assay buffer. Cells were analyzed on the multicolor DNA damage protocol using the Guava Muse® cell analyzer (Millipore).

### Detection of Annexin-V Binding

PEO1 cells and PEO4 cells were treated with 2 µM or 4 µM auranofin for 72 hours. The treated cells were collected and resuspended in different volumes of media to have 1 x 10^5^ to 5 x 10^5^ cells per mL. A 100 µL cell suspension containing approximately 1 x 10^6^ cells were placed in a tube, and 100 µL of the annexin V and dead cell reagent (Luminex) was added for 20 minutes at room temperature in the dark. Annexin V is a calcium-dependent phospholipid binding protein that binds to phosphatidylserine (PS), which translocates to the extracellular surface during early apoptosis. The dead cell reagent detects live and dead cells, by integrating into the membrane of late apoptotic and dead cells due to the loss of membrane structural integrity. The cells were analyzed using the annexin V and dead cell protocol in the Guava Muse® cell analyzer (Millipore).

### Measurement of Caspase-3/7 Activation

PEO1 cells and PEO4 cells treated with 2 or 4 µM of auranofin for 48 hours were collected. A Muse caspase-3/7 kit (Luminex) was used. A 50 µL suspension containing approximately 5 x 10^5^ cells was placed in a tube. Five microliters of the caspase-3/7 reagent working solution, prepared by diluting the caspase-3/7 stock 1:8 with 1 X PBS, was added to the cells and incubated in the dark for 30 minutes in a 37°C 5% CO_2_ incubator. This reagent is bound to a DNA-binding DEVD peptide substrate that, upon activation of caspase-3/7, is cleaved and then translocates to the nucleus to bind DNA and emit fluorescence. Cells were then stained for 5 minutes at room temperature in the dark with 150 µL of Muse caspase-7-AAD substrate working solution prepared at 1:75 dilution using 1X assay buffer. The 7-AAD is a cell permeable DNA-binding dye that integrates through cells that have lost their membrane structural integrity. Analysis was done using the caspase-3/7 protocol on the Guava Muse® cell analyzer (Millipore).

### Treatment with a Caspase Inhibitor

PEO1 cells and PEO4 cells were pre-treated with 50 µM z-DEVD-fmk for 2 hours (Selleck Chemicals, Houston, TX, USA). This is a specific irreversible caspase-3 inhibitor that also exhibits potent inhibition of caspase-6, caspase-7, caspase-8, and caspase-10. Two or 4 µM auranofin were added for 24 hours to PEO1 cells, and for 48 hours to PEO4 cells, and cell viability was assessed by cytometry and analyzed using the Guava Muse® cell analyzer (Millipore).

### Detection of Mitochondrial Membrane Depolarization

PEO1 cells and PEO4 cells treated with 2 or 4 µM of auranofin for 24 hours were collected. Cells were resuspended in 500 µL of 1X assay buffer for a final concentration of 5 x 10^5^ cells per mL. One hundred microliters of the cell suspension were placed in a 1.5 mL centrifuge tube and were incubated for 20 minutes at 37°C 5% CO_2_ with 95 µL of mito-potential working solution prepared by diluting the Muse mito-potential dye at 1:1000 in 1X assay buffer. Five microliters of the Muse mito-potential 7-AAD reagent (Luminex) were added to each tube and incubated for 5 minutes at room temperature. Analysis was done using the mito-potential protocol on the Guava Muse® cell analyzer (Millipore).

### Assessment of Intracellular ROS Levels

To assess whether auranofin stimulates the production of ROS in HGSOC cells, PEO1 cells and PEO4 cells were treated with 8 µM AF for 4 hours. An oxidative stress assay (Luminex) was used to measure the levels of intracellular superoxide. This assay uses a cell permeable reagent named dihydroethidium (DHE) that upon interaction with superoxide ions binds to DNA and produces red fluorescence. Following treatment, the cells were collected and prepared in 1X assay buffer at 1 x 10^6^ to 1 x 10^7^ cells per mL. An intermediate solution of the Muse oxidative stress reagent was prepared by diluting it 1:100 with 1X assay buffer. To prepare the Muse oxidative stress working solution, the intermediate solution was diluted 1:80. One hundred and ninety microliters of the Muse oxidative stress working solution was added to 10 µL of cells and mixed thoroughly by pipetting up and down. The samples were incubated for 30 minutes at 37°C. The stained samples were analyzed using the oxidative stress protocol on the Muse® cell analyzer (Millipore).

### Drug Interaction Between Auranofin and L-BSO

Two hundred thousand PEO1 cells and PEO4 cells were plated per well in 6-well plates and allowed to attach overnight. The cells were treated with 2 or 4 µM auranofin with or without 5 µM of L-BSO for 72 hours. The cells were then collected, and the viability and cell number were measured using the Muse® count and viability reagent. The experiment was repeated three times. The combination index (CI) was then calculated using the method of Chou and Talalay [32] utilizing the CompuSyn Software (ComboSyn Inc. Paramus, NJ, USA). For a specific drug combination, a CI>1 is considered antagonistic, CI=0 means no drug interaction, whereas CI=1 means additivism, and CI<1 denotes synergism.

### Measurement of TrxR Activity

We used a thioredoxin reductase (TrxR) assay kit purchased from Abcam (Cambridge, MA, USA). In this colorimetric assay, TrxR activity is measured by the reduction of 5,5’- dithiobis (2-nitrobenzoic) acid (DTNB) using NADPH to 5-thio-2-nitrobenzoic acid (TNB^2-^), and absorbance is measured at 412 nanometers. Two million PEO1 cells or PEO4 cells were treated with 1, 2, or 4 µM auranofin for 24 hours. Cells were placed on ice, collected by scraping, and washed twice with 1 X PBS. Cells were centrifuged at 1,500 x *g* for 6 minutes, and the supernatant was removed. Cells were resuspended again in 1 mL of 1X PBS and centrifuged at 2,000 x *g* for 5 minutes. The supernatant was decanted, and the cell pellet was snap frozen in liquid nitrogen and stored at -80°C until further processing. The cell pellet was homogenized on ice with 150 µL of cold assay buffer containing 1 X protease inhibitor cocktail (Abcam), and centrifuged at 12,000 x *g* for 15 min at 4°C. The protein concentration of the supernatant was quantified using the BCA protein assay (Pierce). Two sets of 50 µg of protein for each sample, and 10 µL of the TrxR positive control, were loaded into a 96 well plate, and the volume was adjusted to 50 µL with TrxR assay buffer. Ten microliters of TrxR inhibitor were added to one set to test the background enzyme activity, and 10 µL of assay buffer was added to the other set to measure total DTNB reduction. A standard curve was generated with 0, 10, 20, 30, 40 and 50 nmol/well that was adjusted to a final volume of 100 µL with assay buffer. A reaction mix containing TrxR assay buffer, DTNB solution and NADPH was prepared, and 40 µL of the reaction mix was added to the positive control and to each sample and mixed well. The optical density (OD) was measured at 412 nanometers (nm) to obtain A_1AB_ and A_1INH_, and the samples were incubated for 20 minutes at 25°C and measured at 412 nm again to obtain A_2AB_ and A_2INH._ To determine the optical density of TNB^2-^ generated by TrxR, the following calculation was used: ΔA_412nm_ = (A_2AB_ – A_2INH_) – (A_1AB_ – A_1INH_); AB is assay buffer and INH is inhibitor. The TrxR activity was determined using the following formula: TrxR activity = ΔB/[(T2-T1) x V] x sample dilution factor = nmol/min/mL = mU/mL; ΔB: nmol is calculated by applying ΔA_412nm_ to the TNB standard curve; T_1:_ time of the first reading (min); T_2:_ time of the second reading (min); and V: pretreated sample volume (mL). One unit of TrxR is the amount of enzyme that generates 1.0 µmol of TNB per minute at 25 °C.

### In Vitro Analysis of Total GSH

A GSH assay kit was purchased from Abcam. This colorimetric assay measures the concentration of reduced GSH *in vitro.* The kit contains a chromophore, in which the reduction of the chromophore by an enzyme can be measured kinetically by the absorbance at 450 nanometers. The absorbance is directly proportional to the amount of GSH that is present in each sample. The PEO1 cells and PEO4 cells were treated with 1 or 2 µM auranofin in the presence or absence of 5 µM L-BSO for 24 hours. Cells were placed on ice, collected by scraping, and washed twice with 1 X PBS. Cells were centrifuged at 1,500 x *g* for 6 minutes, and the supernatants were removed. Cells were resuspended again in 1 mL of 1X PBS, split in two tubes to have two sets of samples, and centrifuged at 2,000 x *g* for 5 minutes. The supernatants were decanted, and the cell pellets were snap frozen in liquid nitrogen and stored at -80°C until further processing. One set of samples was used to determine the protein concentration in mg/mL. The other set of samples was homogenized on ice using 100 µL of 5% sulfosalicylic acid, vortexed, and kept on ice for 10 minutes. The samples were then centrifuged at 12,000 x *g* at 4°C for 20 minutes, and the supernatant was collected and kept on ice. The samples were diluted 5-fold using the GSH assay buffer, and 10 µL of the diluted samples were added per well in a 96-well plate for the sample well and the sample background control well. The volume of each sample was adjusted to 20 µL/well with GSH assay buffer. A standard curve was produced with 0, 0.4, 0.8, 1.2, 1.6, and 2 nmol/well of the GSH standard that was adjusted to 20 µL/well with the GSH assay buffer. A reaction mix containing substrate mix A, diluted enzyme mix A, enzyme mix B, enzyme mix C, and substrate mix B was prepared; 80 µL of the reaction mix was added to each sample wells and the GSH standard wells. A background control mix was prepared containing everything except the diluted enzyme mix A, and 80 µL of the background control mix was added to the wells of the sample background controls. The absorbance was then measured at 450 nanometers kinetically for 60 minutes at room temperature, and the absorbances at two time-points within the linear range were selected for each sample. The concentration of GSH was then determined using a formula recommended by the provider as follows: 1) calculate the rate of each standard reading and sample reading; Rate = [(OD_2_ – OD_1_)/t_2_ – t_1_)] where OD_2_ is the optical density for the second time point, and OD_1_ is the optical density for the first time point; t_1_ is the initial time in min and t_2_ is the second time in min. Subtract the 0-standard rate from all the standard rates, and plot the GSH standard curve to obtain the slope of the curve; 3) calculate the rate of the background corrected samples by subtracting the sample background control rate from the sample rate; Rate background corrected samples = [rate _sample_ – rate _background control_]; 4) apply the rate of the background corrected samples to the GSH standard curve to calculate the amount of GSH in each sample: B = Rate background corrected samples /slope of the standard curve; and 5) calculate the GSH amount in sample as (B/ [VxP]) x D = nmol/mg. B is the amount of GSH from the standard curve (nmol); V is the volume of sample added in each well (mL); P is the protein concentration in mg/mL; and D is the sample dilution factor.

### Statistics

For tests involving Western blot analysis, the experiments were repeated at least twice with a similar outcome. All other data represent triplicates and are expressed as the mean ± SEM. Differences were considered significant if p<0.05. GraphPad Prism 9 (GraphPad Software, La Jolla, CA, USA) allowed for statistical analysis of data using *t*-test to compare two groups, or one-way ANOVA followed by Tukey’s multiple comparison test to compare more than two groups within an experiment.

## Results

### Auranofin reduces the vitality of HGSOC cells regardless of their sensitivities to cisplatin

To test whether blocking TrxR impairs the wellbeing of ovarian cancer cells, we exposed sibling cell lines to auranofin and measured their vitality. We used a pair of cell lines termed PEO1 and PEO4 which have different sensitivities to platinum. For instance, we recently demonstrated that PEO4 cells are about ten times less sensitive to cisplatin than their PEO1 siblings obtained from the same patient earlier during disease evolution [15]. Despite the large difference in platinum sensitivity among the two cell lines (Figures 1A and 1B), both cellular types responded to increased concentrations of auranofin with similar impairment in wellbeing as denoted by the similar decrease in their vitality (Figures 2C and 2D).

**Figure 1.**
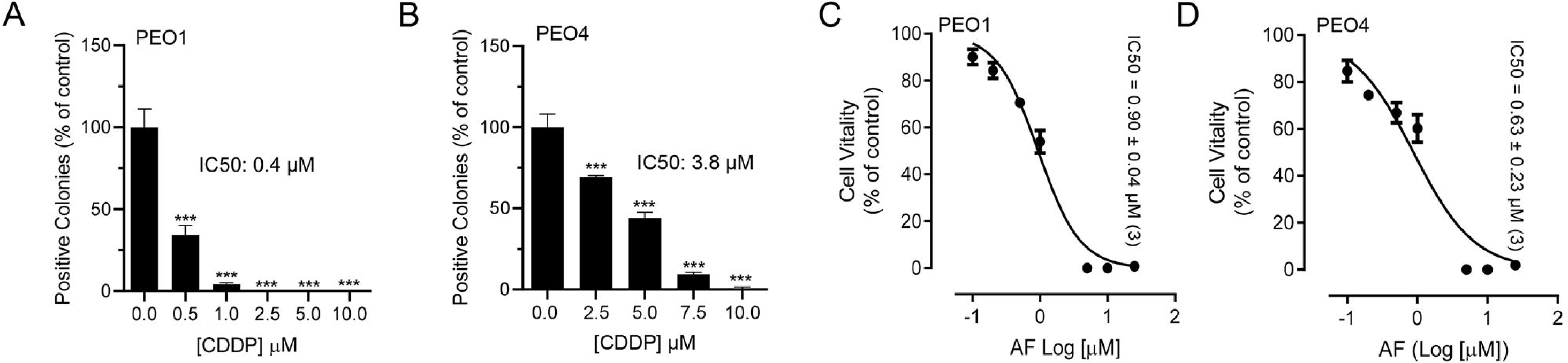
Effect of auranofin (AF) on the vitality of PEO1 or PEO4 cells. Cells were treated with DMSO (vehicle) or with various concentrations of cisplatin (CDDP) or auranofin (AF) for 72 hours. At the end of the treatment, the cells were subjected to a clonogenic survival assay (for CDDP) and vitality assay (for AF) as detailed in materials and methods. Panels A and B show the contrasting clonogenic survival among the cell lines in response to CDDP whereas panels C and D show a similar decrease in vitality caused by AF in the two cell lines.

**Figure 2.**
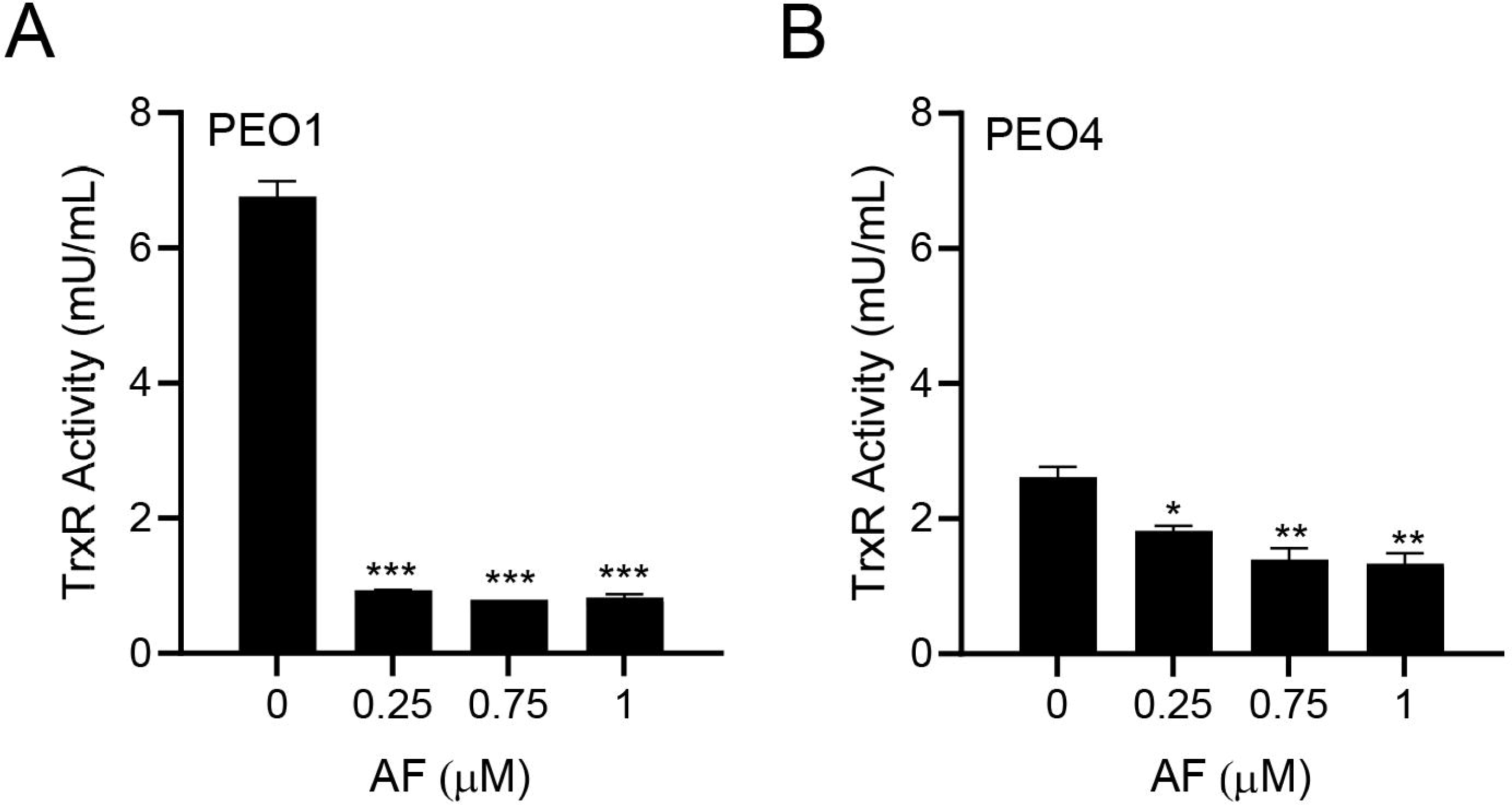
Effect of auranofin (AF) on the activity of the enzyme TrxR. In this colorimetric assay, TrxR activity was measured by the reduction of 5, 5’-dithiobis (2-nitrobenzoic) acid (DTNB) using NADPH, to 5-thio-2-nitrobenzoic acid (TNB^2-^).*p<0.05, **p<0.01, and ***p<0.001 when compared to vehicle.

### Auranofin inhibits TrxR activity

As auranofin is known to target TrxR with an inhibitory effect [22], we tested whether indeed this occurred in PEO1 and PEO4 cells. PEO1 cells had relatively high basal TrxR activity that is comparable to the rat liver homogenate used as positive control (data not shown). Once treated with different concentrations of auranofin for 24 hours, there was a potent inhibition of TrxR activity (Figure 2A). In PEO4 cells, basal TrxR enzymatic activity was lower than that found in PEO1; nevertheless, the activity of TrxR was significantly reduced further by auranofin in a concentration-dependent manner (Figure 2B). Our results demonstrate that auranofin hinders the activity of its primary target, TrxR, in HGSOC cells.

### Auranofin triggers the accumulation of reactive oxygen species

Although TrxR activity was assessed, a direct measure of the influence of inhibiting TrxR is the causation of oxidative stress. Thus, to confirm the effect of auranofin on oxidative stress we measured ROS production in HGSOC cells in the presence or absence of auranofin for 4 hours. Figure 3A and 3B show the significant increase in the percentage of ROS positive cells in response to auranofin in both clinically platinum sensitive PEO1 cells and clinically platinum resistant PEO4 cells.

**Figure 3.**
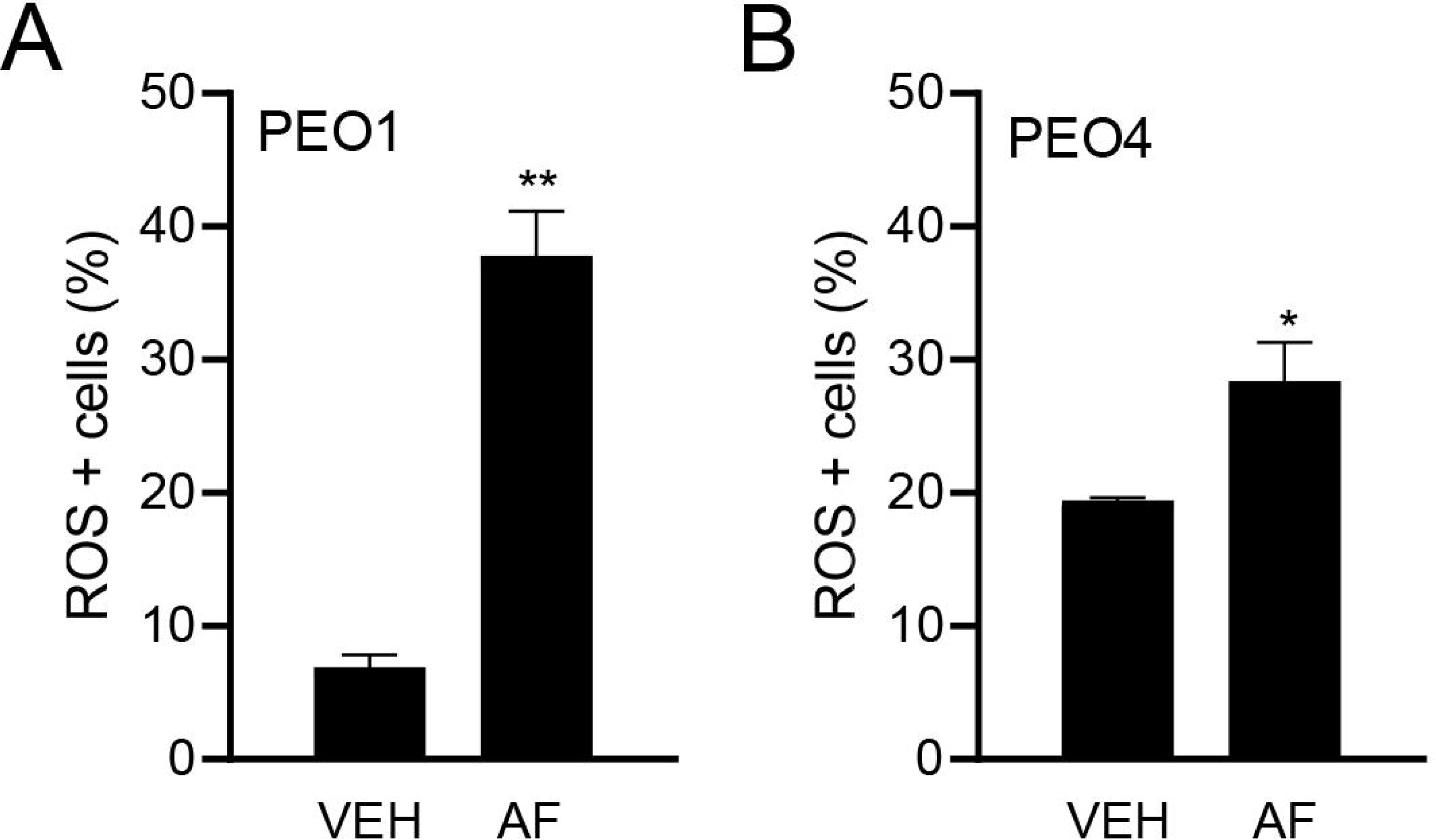
Effect of vehicle (VEH) or auranofin (AF) on the production of ROS. The oxidative stress method utilized measures the levels of a cell permeable reagent named dihydroethidium (DHE) that upon interaction with superoxide binds to DNA and produces a red fluorescence. Treatment was done with 8 μM AF for only 4 hours. *p<0.05 and **p<0.01 compared to vehicle.

### Auranofin causes lethality of HGSOC cells in association with induction of apoptosis

The reduction in the vitality or metabolic activity of cells exposed to auranofin shown in Figure 1 suggests that the drug may have cytotoxic effects. Indeed, auranofin showed reduction in viability (Figure 4A and 4E), which was associated with an increase in markers of apoptotic cell death, such as double labelling of Annexin-V and 7-AAD (Figure 4B and 4F). Confirmation of apoptosis induced by auranofin was elicited by the accumulation of cells with hypodiploid DNA content (Figure 4C and 4G). Finally, if cells that were still viable after 72 hours of treatment (Figures 4A and 4E) were placed in a clonogenic survival plate in the absence of auranofin, the long-term toxicity of the previous exposure to the drug was clearly depicted by the concentration-dependent reduction of viable colonies (Figure 4D and 4H). In contrast to results obtained with vitality, in which both cells seem to respond to auranofin in a similar fashion, measuring lethality-related parameters clearly show that auranofin is more potent against PEO1 cells than to PEO4 cells. For instance, clonogenic survival depicts a fivefold difference, with an IC50 of 0.53 μM for PEO1 cells (Figure 4D) and an IC50 of 2.8 μM for PEO4 cells (Figure 4H).

**Figure 4.**
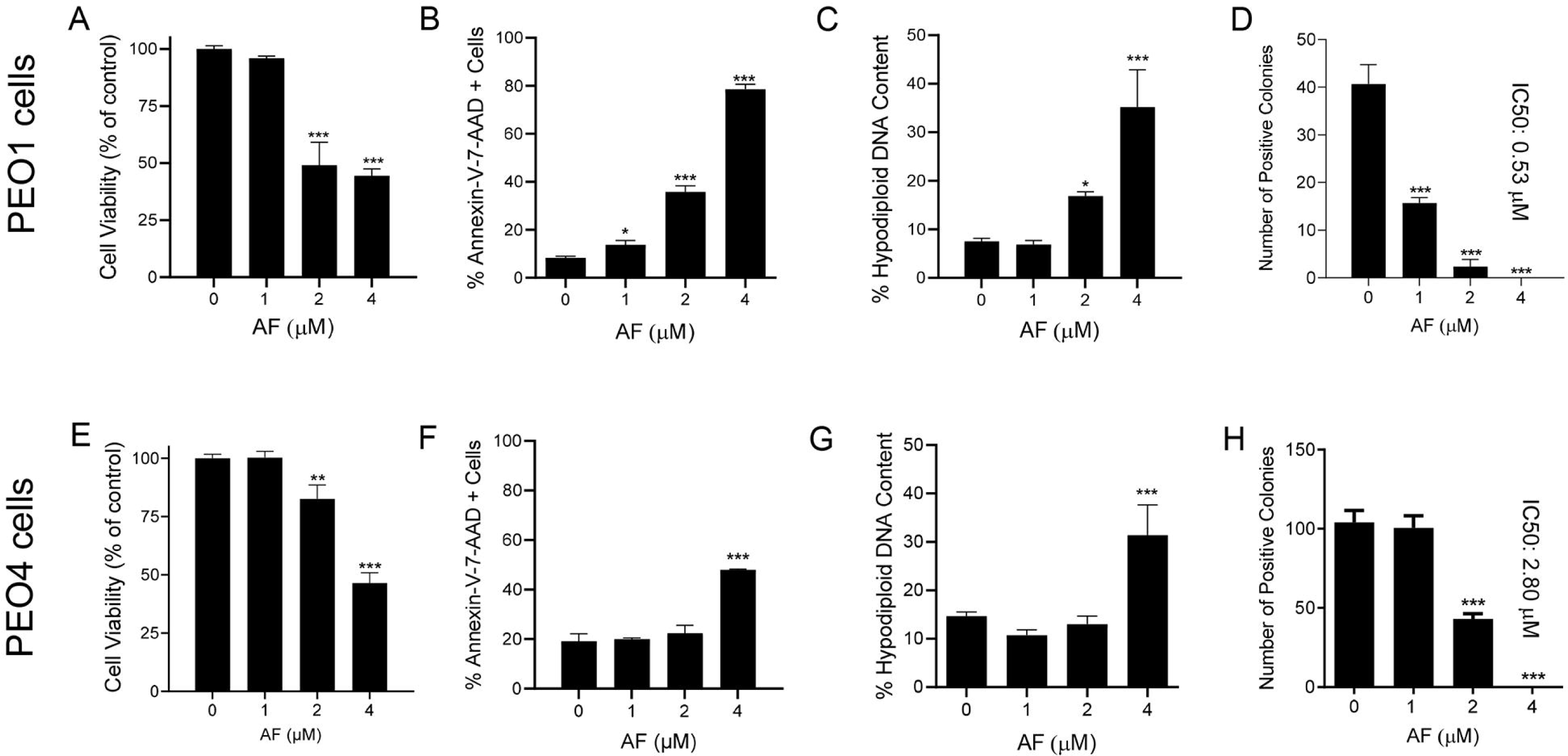
Viability of PEO1 and PEO4 cells after 72 hours of treatment with auranofin (AF) assessed using a cytometric viability assay (A and E). In a similar experiment, cells were stained with Annexin V and 7-AAD to determine apoptosis (B and F). Cells remaining from the viability experiment were also stained with propidium iodide and studied for cell cycle distribution; only the hypodiploid DNA contents are shown (C and G). Finally, cells that were still alive after 72 hours of treatment with AF shown in (A) and (E), were subjected to a clonogenic survival assay to define their long-term reproductive capacity (D and H). * P<0.05, ** P<0.01 and *** P<0.001 compared with cells treated with vehicle.

### Auranofin induces dissipation of the mitochondrial potential, a phenomenon that is prevented by the presence of the ROS scavenger N-acetyl cysteine

Cellular energy produced during mitochondrial respiration is stored as an electrochemical gradient across the mitochondrial membrane, and this accumulation of energy in healthy cells creates a mitochondrial trans-membrane potential (ΔΨ_m_) that enables the cells to drive the synthesis of ATP. Collapse of this potential is believed to coincide with the opening of the mitochondrial permeability transition pores, leading to the release of cytochrome C into the cytosol, which then triggers the downstream events in the apoptotic cascade [33,34]. Depolarization of the inner mitochondrial membrane potential is thus a reliable indicator of mitochondrial dysfunction and cellular death by apoptosis [35]. We here show that treatment of PEO1 cells or PEO4 cells with auranofin caused the loss of ΔΨ_m_and further illustrate that the presence of the ROS scavenger N-acetyl cysteine (NAC) [36] prevented the depolarization of the mitochondrial membrane (Figure 5A and 5B).

**Figure 5.**
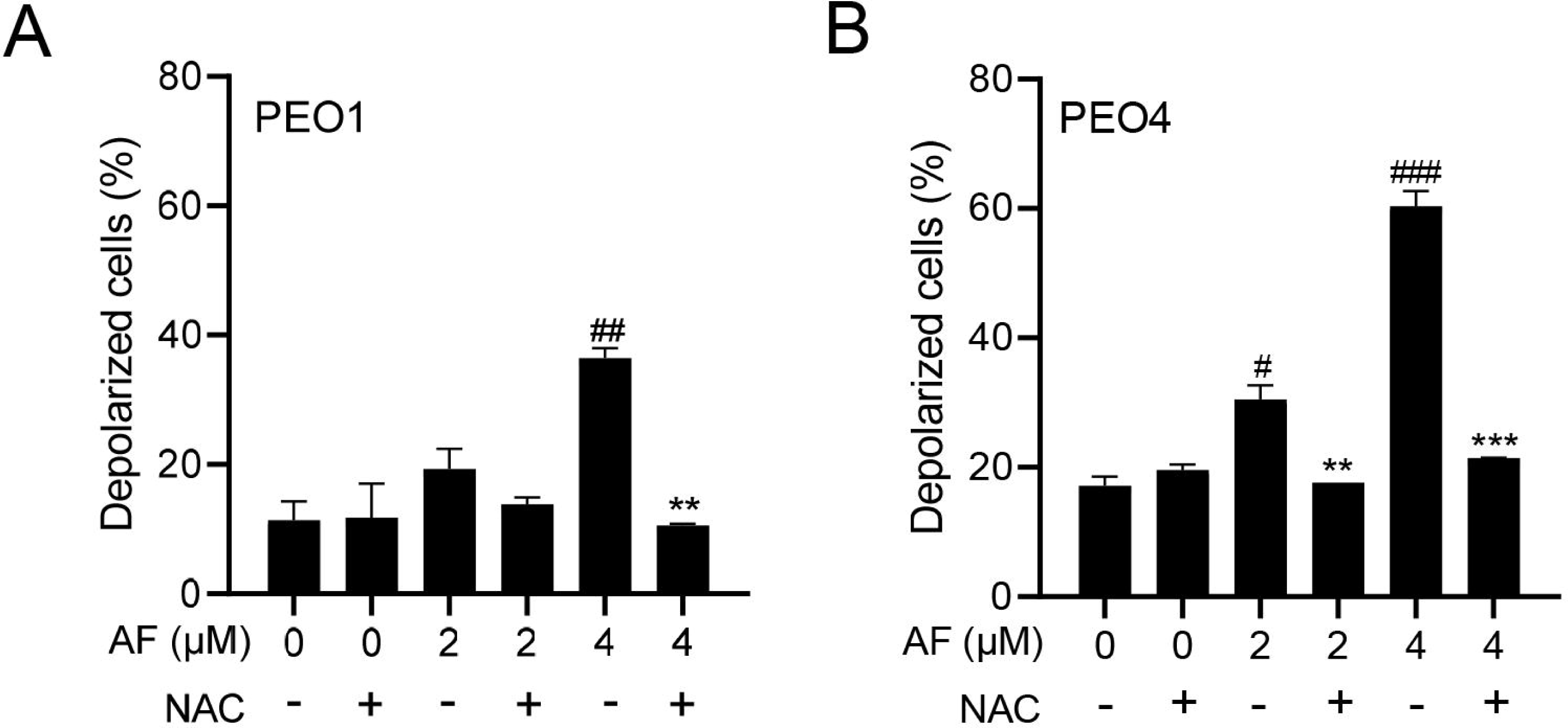
Panels show the percentage of cells with depolarized mitochondria that increases with auranofin (AF). # P<0.05, ## P<0.01 and ### P <0.001 when compared to vehicle treated controls; ** P<0.01 and *** P<0.001 when compared to the corresponding AF-treated groups. Mitochondrial depolarization was assessed using a cytometric method that utilizes a mito-potential dye. High membrane potential drives accumulation of mito-potential dye within the inner membrane of intact mitochondria resulting in high fluorescence. Cells with depolarized mitochondria demonstrate a decrease in fluorescence. NAC, N-acetyl cysteine.

### Auranofin-induced cell death is prevented by N-acetyl cysteine

To determine whether the mechanism of cytotoxicity by auranofin is dependent on the production of ROS, PEO1 cells or PEO4 cells were cultured in the presence of 2 or 4 μM auranofin with the addition or not of 5 mM NAC. Results presented in Figure 6 show that auranofin reduced the viability in a concentration-related manner. However, the presence of the antioxidant NAC could prevent the lethality of auranofin (left panels in Figure 6). The right panels in Figure 6 clearly show the morphological deterioration of the cell cultures in the presence of auranofin and how in the presence of NAC they resemble back the cellular morphology of vehicle-treated cells.

**Figure 6.**
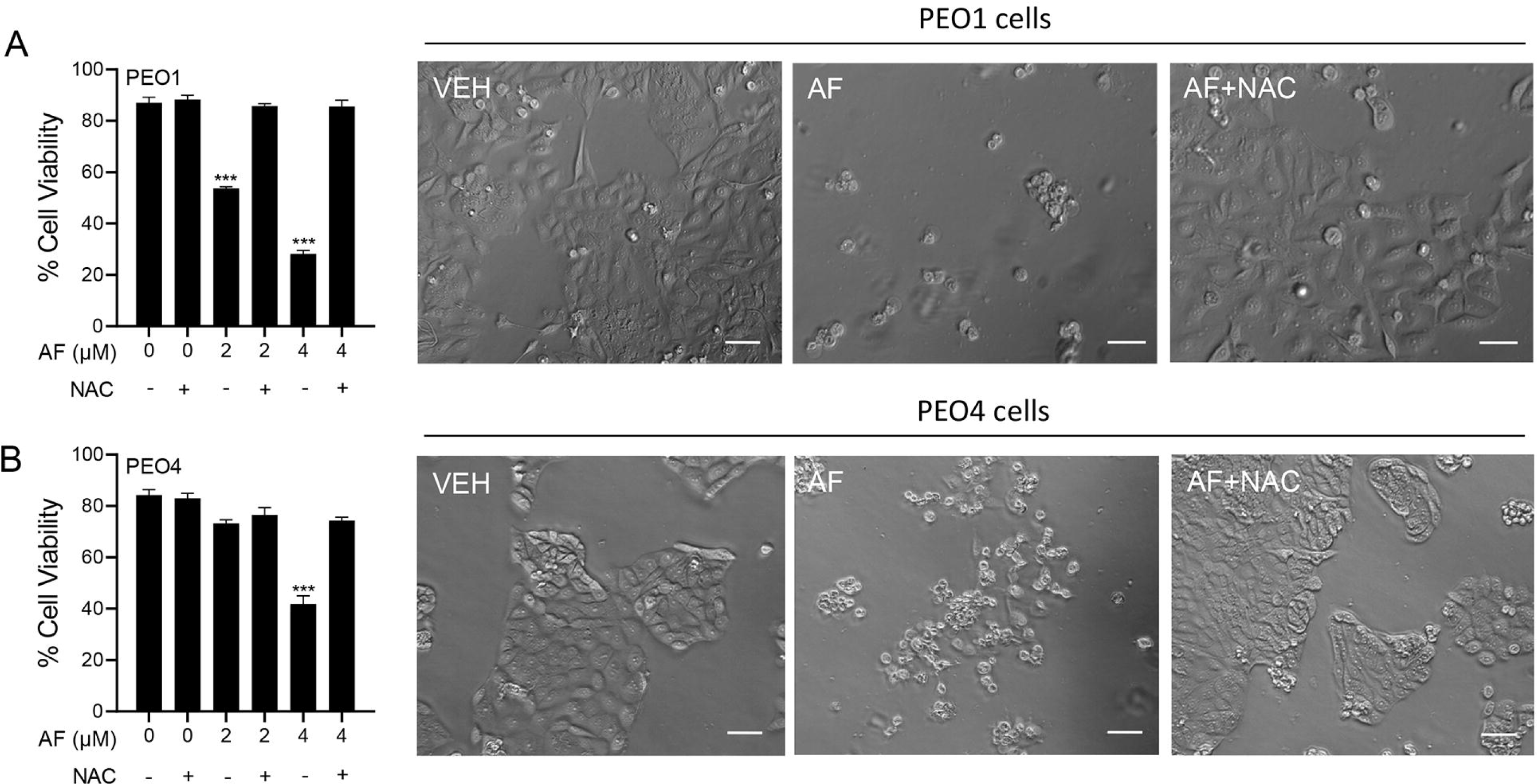
PEO1 cells (A) or PEO4 cells (B) were treated with auranofin (AF) in the absence or presence of 5 mM NAC. Cell viability was assessed after 72 hours by microcytometry. *** P<0.001 vs. AF+NAC. In the right panels, phase contrast images were obtained after 72 hours of incubation with the indicated drugs. VEH, vehicle; AF, auranofin; NAC, N-acetyl cysteine. Scale bar = 50 μm.

### NAC prevented auranofin-induced caspase-3/7 activation, cleavage of PARP, and induction of γH2AX

A caspase-3/7 cytometric assay was utilized to define the activation of the executer caspases in response to auranofin. In this assay, it was clear that auranofin was more potent in inducing caspase-3/7 activation in PEO1 cells than in PEO4 cells; nonetheless, such activation was blunted by the presence of NAC (Figure 7A and 7C). Likewise, and in both cell lines, when assessing induction of apoptosis by measuring the cleavage of PARP, we observed that auranofin was effective in inducing such a cleavage, which was however, at least in part, prevented by NAC (Figure 7B and 7D). Finally, auranofin triggers the accumulation of the DNA damage marker γH2AX in both PEO1 (Figure 7E) and PEO4 (Figure 7F). Such accumulation of γH2AX was entirely abrogated by NAC. These results suggest that both executer caspase activation and DNA damage are mediated by generation of reactive oxygen species (ROS).

**Figure 7.**
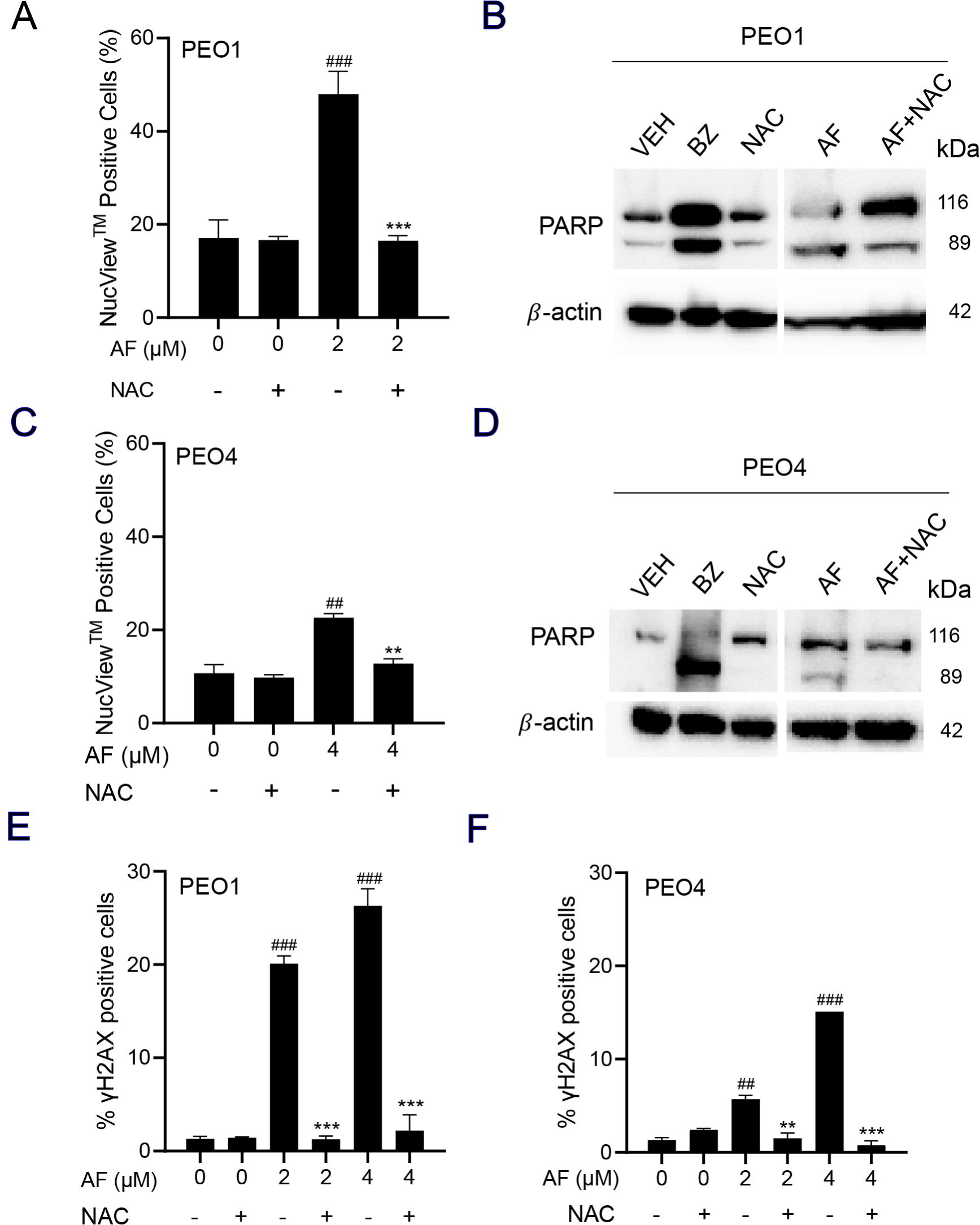
PEO1 cells (A) or PEO4 cells (B) were treated with auranofin (AF) in the absence or presence of 5 mM NAC for 24 hours (to measure PARP), 48 hours (to measure caspase-3/7 activity) or 72 hours (to measure accumulation of γH2AX). In A and C, cells were exposed to the Muse® Caspase-3/7 reagent, which is a cell membrane permeable in combination with a dead cell dye (7-AAD). ## P<0.01 and ###P<0.001 compared to vehicle. **P<0.01 and ***P<0.001 compared to AF. (B) and (D) depict the effect of AF and NAC on the cleavage of PARP as detected by western blot. In this experiment, cells treated with 20 nM bortezomib (BZ) were used as a positive control of PARP cleavage. E and F show the accumulation of γH2AX in response to AF with and without NAC. ## p<0.01 and ###p<0.01 compared to vehicle. ** p<0.01 and ***p<0.001 compared to the respective concentration of AF.

### The cytotoxic effect of auranofin and L-BSO against HGSOC is synergistic and associates with a further increase in ROS production and reduced levels of GSH

We hypothesized that the toxicity of auranofin could be augmented by the blockage of an additional, likely compensatory antioxidant system: GSH. Thus, we decided to simultaneously block TrxR with auranofin and GSH with L-BSO [37]. In Figure 2, we show that auranofin blunts TrxR activity. In Figure 8, we show that the combination of auranofin and L-BSO lead to a further reduced viability when compared to auranofin monotherapy (Figure 8A and 8E). The interaction between auranofin and L-BSO is synergistic based on the CI method of Chow and Talalay (Figure 8B and 8F). Furthermore, the presence of auranofin, as anticipated earlier, increased ROS levels in both cell types, yet such elevation was furthered by the combined presence of L-BSO (Figure 8C and 8G). Finally, we show that blockage of TrxR by auranofin treatment alone leads to a compensatory increase in GSH, which is however significant reduced by the presence of L-BSO (Figure 8D and 8H).

**Figure 8.**
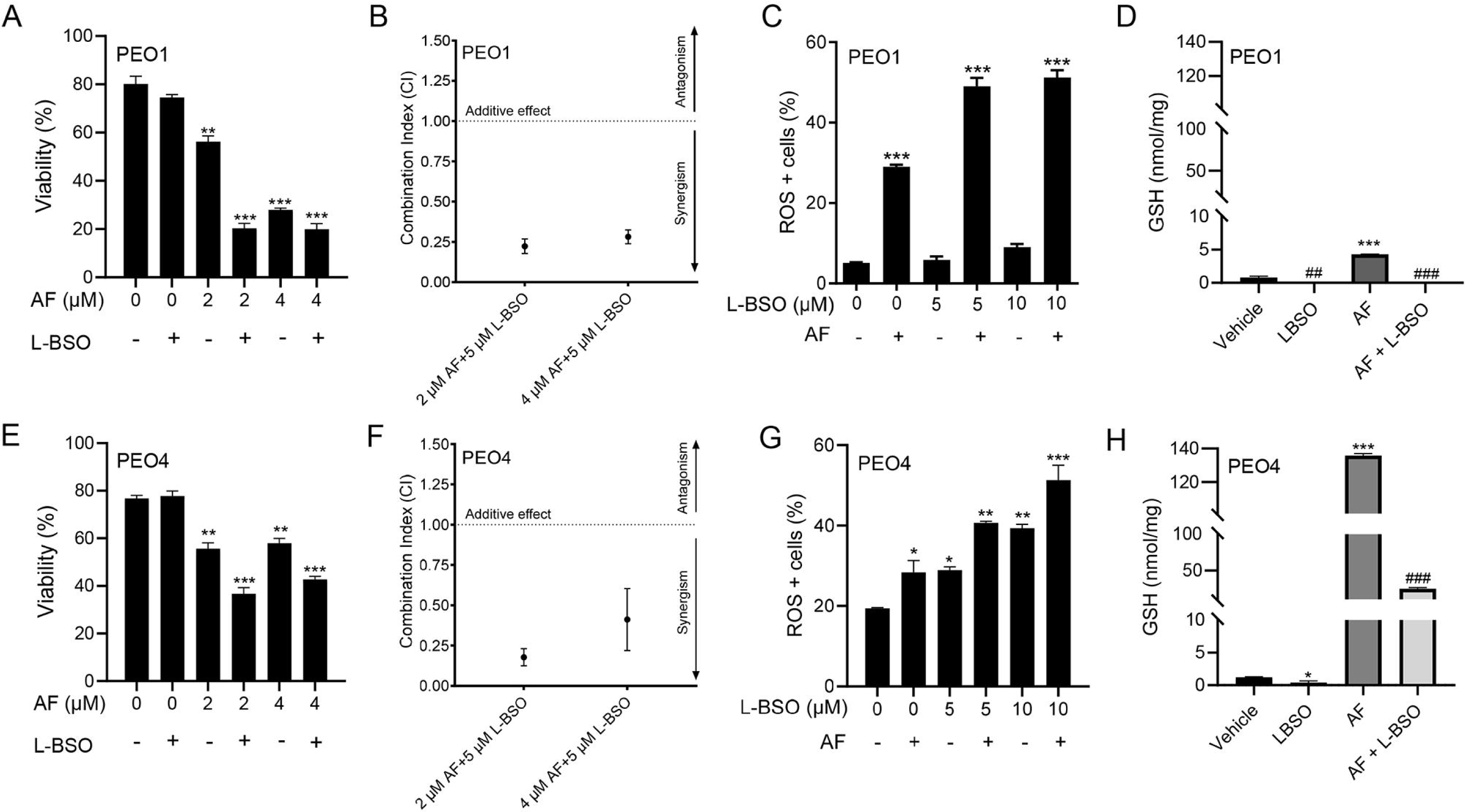
PEO1 cells (A) or PEO4 cells (E) were treated with 2 or 4 μM auranofin (AF) for 72 hours in the presence (+) or absence (-) of L-buthionine sulfoximine (L-BSO) and viability was recorded by cytometry. (B) and (F) The combination indexes of the drug interaction using various combinations of AF and L-BSO are shown. All intersections show a CI <1 indicating synergism among the drugs. The CI were calculated using the viability data of cells treated with the depicted concentrations of AF and/or L-BSO in three independent experiments. (C) and (G) show ROS levels upon action of AF with (+) or without (-) L-BSO. (D) and (H) display GSH levels respectively in PEO1 cells or in PEO4 cells treated with vehicle, 2 μM AF, or the combination of 2 μM AF and 5 μM L-BSO. In (A), (C), (E), and (G), **p<0.01 and ***p<0.001 vs. control. In (D) and (H), ***p<0.001 vs. vehicle, #p<0.05 and ##p<0.01 vs. vehicle, and ###p<0.001 vs. AF.

### The lethal effect of auranofin and L-BSO against HGSOC is prevented by the presence of the ROS scavenger NAC

We studied whether the cytotoxicity of auranofin in combination with L-BSO was prevented by the ROS scavenger and antioxidant NAC. In Figure 9A for PEO1 cells and Figure 9C for PEO4 cells, we show that the toxicity caused by 2 μM auranofin to the cells was greatly enhanced by the presence of 5 μM L-BSO. Of interest, such reduction in viability was almost totally reversed by the presence of NAC. The phase contrast panels in Figure 9B and 9D show that the morphology and growth of the cells was negatively affected by auranofin alone and furthered by auranofin when in combination with L-BSO. Consistent with the viability data shown in panels A and C, the morphology of cells receiving auranofin and L-BSO was highly affected in a negative manner, but in the presence of NAC, it resembles back to that of vehicle-treated controls.

**Figure 9.**
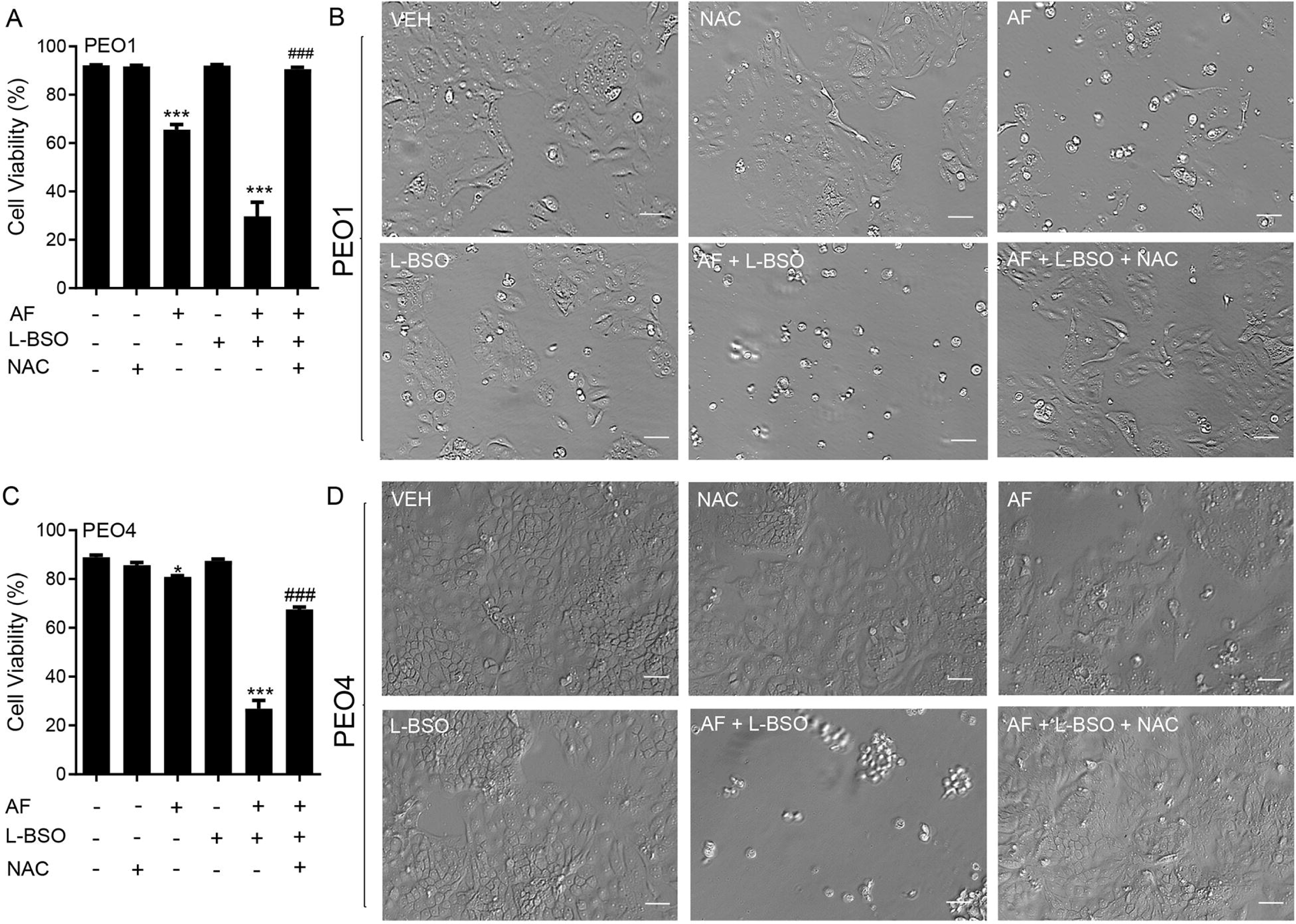
Cell viability (A and C) and phase-contrast images (B and D) of PEO1 and PEO4 cells receiving vehicle (VEH), N-acetyl cysteine (NAC; 5 mM), auranofin (AF), L-buthionine sulfoximine (L-BSO; 5 μM), or the combination of AF/L-BSO or AF/L-BSO/NAC. *P<0.05 and ***P<0.001 compared to vehicle; ###P<0.001 compared to the AF/L-BSO group. Scale bar = 50 μm.

## Discussion

Mechanisms of cytotoxicity of auranofin as a monotherapy or in combination with other drugs have been studied in various cancers including lung [38-41], breast [42-44], pancreatic adenocarcinoma [25], colorectal [45], gastric [46], mesothelioma [47], melanoma cells [48], malignant B-cells and acute lymphoblastic leukaemia (ALL) [49-51]. In ovarian cancer, auranofin has been shown to be efficient in blocking the growth of A2780, SKOV-3, OVCAR-5, IGROV-1, and OV2008 cells [20,27,52-54]. However, despite popular, all these cell lines are unlikely to represent the most common histotype of ovarian cancer we study in this manuscript: high-grade serous ovarian cancer [55]. In our study, we used two HGSOC cell types, PEO1 (platinum sensitive) and PEO4 (platinum resistant), which have been isolated from the same patient throughout the course of the disease [29] and have been genotypically identified as HGSOC [30]. These cell types provide a representative model of the heterogeneous clinical pathological characteristics of the disease, in which PEO1 cells contain the *BRCA2* germline mutation, whereas PEO4 cells have the restored version of the gene [30,56]. Of interest, in a work that studied the combination of auranofin with HSP90 inhibitors against PEO4 cells, it was reported that these cells were resistant to the combination approach [57]. Our results disagree, as we show cytotoxicity by auranofin alone or in combination with the GSH inhibitor L-BSO against both the platinum sensitive PEO1 cells and their sibling platinum resistant PEO4 cells.

One of the advantages of using auranofin as a monotherapy against HGSOC is that the gold complex has been already approved by the FDA against rheumatoid arthritis, and it is currently enrolled in several clinical trials as a monotherapy and in combination with other drugs (reviewed in [22]). This indicates the feasibility of repurposing auranofin against HGSOC as it has been shown to be clinically tolerable.

Our first study that was done to assess the effect of auranofin on the vitality or physiological capability (i.e., wellbeing, as defined in yeast [31]), revealed that both PEO1 and PEO4 cell types were equally affected by the drug. In the context of cell vitality, this means that the physiological wellbeing of PEO1 cells and PEO4 cells—assessed through the surrogate activation of mitochondrial enzymes—was impaired equally in both cell types by auranofin and that there is no cross resistance between the platinum agent and the gold complex. However, when we studied viability (i.e., the capacity of auranofin to kill the cancer cells), we observed that the drug was more efficacious against PEO1 cells than towards PEO4 cells, suggesting that in terms of lethality there is cross resistance between platinum and auranofin. Furthermore, other short-term cytotoxic responses to stress by auranofin that we studied showed that PEO4 cells were again less sensitive to the drug than the PEO1 cells. First, the induction of late apoptosis/death by auranofin, which was represented by positive staining of Annexin V and 7- AAD show a stronger response in PEO1 than in PEO4 cells. The induction of apoptosis by auranofin as one of the principal short-term cytotoxic effects was shown also in multiple myeloma, acute myeloid leukemia, murine triple-negative breast cancer, lung cancer, and mesothelioma cells [23,38,49,58]. We confirmed the induction of apoptosis with the accumulation of hypo-diploid DNA content during the short-term exposure to auranofin, which coincides with the increase in the sub-G1 DNA content elicited by auranofin in an experiment done with non-small cell lung carcinoma (NSCLC) cells [59]. The short-term cytotoxic effects of auranofin started to be observed at a concentration of 2 µM in PEO1 cells, except for annexin V/7-AAD staining which shows positivity already in response to 1 µM of the drug. The colony formation assay showed that auranofin, even in those cells that were not killed in a short term, elicits long-term inhibition of their reproductive ability by inhibiting their clonogenic survival capacity; such reduction in clonogenicity was again more pronounced (∼ five folds) in PEO1 cells than in PEO4 cells. Inhibition of positive colony formation by auranofin was also demonstrated in stem-like cancer cell side population and in SKOV-3 epithelial ovarian cancer cells in a p53-independent manner [60,61].

When we explored the direct effect of auranofin against TrxR activity, the presumed target of the drug when acting against rheumatoid arthritis [17], we observed a marked inhibition of the activity as expected. However, this inhibition was achieved using much lower concentrations than the ones needed for achieving a cytotoxic effect, suggesting that the inhibition of the activity of TrxR is not sufficient to kill HGSOC cells. Similarly, TrxR activity was inhibited by auranofin at non-cytotoxic doses in Calu-6 lung cancer cells [38]. There are other studies, however, that show that auranofin inhibits TrxR activity in various cancers at cytotoxic concentrations, including the non-HGSOC ovarian cancer cell line A2780 [38,58,62-64]. Although TrxR activity was inhibited by non-lethal doses of auranofin in both PEO1 cells and PEO4 cells, the inhibition was less significant in PEO4 cells than in PEO1 cells primarily due to the lower level of basal TrxR activity displayed by PEO4 cells when compared to PEO1 cells. This may explain the apparent lower sensitivity to the cytotoxicity of auranofin displayed by PEO4 cells, which may rely less on the TrxR antioxidant system for survival when compared to PEO1 cells; such higher sensitivity of PEO1 cells versus PEO4 cells to the toxicity of auranofin was demonstrated in parameters such as the reduction of cell viability, the reduction in positive colony formation, the increase in annexin V/7-AAD binding, and the increase in hypodiploid DNA content.

The inhibition of the TrxR enzymatic activity by other compounds has consequently resulted in increased oxidative stress [65]. Confirming that inhibition of TrxR activity by auranofin in HGSOC is also associated with elevated oxidative stress, we observed elevation of ROS production in PEO1 cells and PEO4 cells following exposure to the gold complex. The increase in ROS levels was more evident in PEO1 cells but the basal levels of ROS were instead higher in the platinum resistant sibling PEO4 cells. This observation is in agreement with findings in the literature claiming that high basal expression of ROS associates with increased resistance to platinum-based chemotherapy [66]. Such elevated levels of ROS in platinum resistant cells are tolerated by the expression of antioxidant genes resulting from the interaction of mutated p53 with the ROS-sensitive transcription factor nuclear factor erythroid 2-related factor 2 (NRF2) [67,68]. This indicates that mutations in p53 as observed in HGSOC can lead to increased ROS levels with a compensatory elevation of antioxidant activity for protection [66].

Once we confirmed that the production of ROS in HGSOC is induced by auranofin, we decided to explore whether ROS plays a role in the cytotoxic effects that are elicited by the gold complex. We used N-acetyl cysteine (NAC), an agent that reduces ROS indirectly via the upregulation of NRF2 and increased synthesis of the antioxidant, glutathione (GSH), by providing cysteines [36]. In a direct manner, NAC can scavenge ROS molecules when it is metabolized into sulfane sulfur species thus exhibiting a cytoprotective role [69]. We found that the presence of NAC reverses the lethality and the detrimental morphological effects that are induced by auranofin in PEO1 cells and in PEO4 cells, demonstrating that these cytotoxic effects are primarily dependent on ROS-induced damage. Of interest, in PEO4 cells, the reversal of auranofin-induced cell death by NAC is only observed when challenged with 4 µM of the gold complex, indicating that the clinically platinum resistant PEO4 cells are less sensitive to ROS-induced cytotoxicity by auranofin than the clinically platinum sensitive PEO1 cells. Although there is a difference in sensitivity to ROS induction by auranofin in the two cell types, the protection by NAC in both cell lines emphasizes the important role of maintaining moderate ROS levels for promoting cell proliferation in HGSOC.

In most parameters of cytotoxicity that we assessed, PEO4 cells were less sensitive to auranofin than PEO1 cells. Interestingly, however, when analyzing the mitochondrial membrane potential in response to auranofin, PEO4 cells showed more sensitivity to auranofin than the PEO1 cells did. This is a significant finding because the mitochondrial membrane potential plays a critical role in whether the cell undergoes death as part of the intrinsic apoptotic pathway [70]. A study on the mitochondrial activity in platinum-resistant NSCLC cells showed that increased mitochondrial mass is associated with cisplatin resistance and increased cell proliferation in A549 lung cancer cells [71]. The study done by Gao et al. on mitochondrial activity reveals that cisplatin-resistant A549 lung cancer cells are highly dependent on mitochondrial function [71], making the mitochondria an attractive therapeutic target. Similarly, a study done on cisplatin-sensitive OV2008 and cisplatin-resistant OV2008 C13* ovarian cancer cells showed that the resistant OV2008 C13* cells had increased mitochondrial fusion, due to better mitochondrial activity, than that seen in the cisplatin-sensitive OV2008 cells [72]. On the other hand, a study exploring the effect of cisplatin on the mitochondria of ovarian cancer cells reveals that human ovarian cancer cells that are sensitive to cisplatin contain increased mitochondrial content in comparison to ovarian cancer cells that are resistant to cisplatin [73]. Another study claims that non-HGSOC cisplatin-resistant OV2008 C13* ovarian cancer cells elicit reduced mitochondrial function in comparison to their cisplatin-sensitive OV2008 sister cell line [74]. These findings suggest that there is a controversy that remains to be addressed on whether cisplatin sensitivity is directly linked to mitochondrial function. However, in our case, it seems that PEO4 cells may have higher mitochondrial function due to their increased sensitivity to the depolarization of the mitochondrial membrane by auranofin in comparison to PEO1 cells. In addition, the depolarization of the mitochondrial membrane by auranofin in PEO1 cells and in PEO4 cells was dependent on the production of ROS. This coincides with the events that occur during the normal metabolic process of the cell, in which ROS is released as a by-product of oxidative phosphorylation to generate ATP [75]. When ROS is released in the mitochondria, it results in the rapid depolarization of the mitochondrial membrane [75].

In addition to the disruption of the mitochondrial membrane potential by auranofin, we also found an increase in the activation of the executor caspases-3/7 and cleavage of PARP induced by auranofin in both PEO1 cells and PEO4 cells. However, the activation of caspase-3/7 and cleavage of PARP by auranofin was higher in the platinum sensitive PEO1 cells than in the platinum resistant PEO4 cells. Activation of caspase-3/7 by auranofin was also reported in mutant p53 NSCLC cells [18]. Other studies demonstrated that auranofin induced caspase 3- mediated apoptosis in p53-null SKOV3 ovarian cancer cells [61], caspase-3 activation and PARP cleavage in human gastric cancer cells [46] and in chronic lymphocytic leukemia (CLL) [76]. Although caspase-3/7 were activated in the PEO1 cells and PEO4 cells by auranofin, cell death induced by auranofin in HGSOC was not dependent on the activation of executer caspases as the decrease in viability was not prevented by the presence of a pan-caspase inhibitor (Figure S1). Interestingly, increased caspase-3/7 activity and PARP cleavage that was induced by auranofin in PEO1 cells and PEO4 cells was dependent on ROS production as it was blocked by NAC. Similarly, cleavage of caspase-3 and PARP by auranofin is ROS-dependent in A549 human lung cancer cells [38]. Likewise in human gastric cancer cells, caspase-3 cleavage by auranofin was reversed in the presence of the ROS scavenger NAC [46], whereas PARP cleavage by auranofin was reversed in the presence of NAC in CLL [76].

Apoptosis commonly occurs as a secondary response to sufficient DNA damage to prevent the survival of cells that contain genomic instability [77]. Since we found that auranofin elicits caspase-3/7-associated apoptosis in HGSOC, we then explored the occurrence of DNA damage. We detected an increased DNA damage response in HGSOC, more significantly in PEO1 cells than in PEO4 cells, following a short-term exposure to auranofin. PEO1 cells were more sensitive to the DNA damage induced by auranofin, likely due to their defective homologous recombination DNA repair machinery and germline mutation in *BRCA2* [30]. Accordingly, the decreased sensitivity of PEO4 cells to DNA damage may be primarily due to their functional DNA repair machinery as a consequence of the reversion of the *BRCA2* mutation [78]. This secondary mutation in *BRCA2* is acquired by various *BRCA2*-mutated cancers as a mechanism of resistance against cisplatin [79]. This indicates that clinically platinum-resistant cells contain a proficient DNA damage response to maintain genome stability and allow the cancer cells to proliferate. Like other cytotoxic mediators induced by auranofin in HGSOC, we found that the DNA damage that is induced by the gold complex in PEO1 cells and in PEO4 cells was ROS dependent, as the antioxidant NAC reversed it. A similar finding was observed in acute lymphoblastic leukemia (ALL), in which auranofin induces DNA damage by increasing ROS levels [49].

Aside from inducing mitochondrial membrane potential loss, DNA damage, and apoptosis, auranofin has been proposed to be a proteasomal deubiquitinase inhibitor [17,42,80]. The proteasome degradation cycle is heavily used by cancer cells to regulate protein degradation and homeostasis, making this pathway an attractive therapeutic target in cancer [81]. In the proteasome degradation cycle, proteins that are meant for degradation are tagged with ubiquitin [82]. Interestingly, we detected an accumulation of poly-ubiquitinated proteins in PEO1 cells and PEO4 cells following treatment with auranofin and demonstrated that such ubiquitination is blocked by NAC, indicating that auranofin-induced poly-ubiquitination is dependent on ROS production (Figure S2). It is unknown from our results, however, whether the poly-ubiquitination is consequence of proteasomal inhibition by auranofin, or, as reported by others, because of inhibition of a deubiquitinase enzyme [28].

Although we have found that auranofin induces cytotoxicity in multiple ways in HGSOC, PEO1 cells showed more sensitivity to the ROS-mediated cytotoxic effects of auranofin than PEO4 cells did. This suggests that auranofin as a monotherapy may not be sufficient for treating recurrent stages of HGSOC that associate with platinum resistance. Auranofin has been shown to elicit anti-cancer effects in various combination treatments, which have been studied in human lung cancer, malignant B cells, breast cancer, gastric cancer, non-HGSOC ovarian cancer, brain tumor cells, and NSCLC (revisited in [22]). However, there have been minimal studies combining auranofin with another agent to target HGSOC cells, the most prevalent histotype of epithelial ovarian cancer [8]. To develop a proficient consolidation therapy against HGSOC, we decided to combine auranofin with another pro-oxidant: L-buthionine sulfoximine (L-BSO). L-BSO is an inhibitor of the rate-limiting enzyme, y-glutamylcysteine synthetase, which is involved in the synthesis of glutathione (GSH) [83]. At moderate levels, GSH plays protective roles within the cell, which includes the removal of ROS, regulation of the cell cycle, and regulation of apoptosis and necrosis [84]. Elevated levels of GSH have been detected in various cancers including breast, ovarian, and lung cancer [85]. Increased levels of GSH are associated with tumor progression and increased drug resistance, primarily due to the maintained levels of oxidative stress. Additionally, elevated levels of GSH have been associated with increased resistance to cisplatin via the increased efflux of cisplatin and redox regulation within the cell [86]. The basal levels of GSH in PEO1 cells and PEO4 cells is controversial. Interestingly, GSH is an attractive therapeutic target in clinically platinum-resistant cancer cells. Thus, our rationale for combining AF with L-BSO against HGSOC is the dual targeting of the antioxidant systems, TrxR and GSH, to combat the recurrent stages of this disease. Multiple studies explored the combination of auranofin and L-BSO in various diseases including mesothelioma cells, human lung cancer, rhabdomyosarcoma, and pancreatic cancer [23,38,63,87]. However, there have been no studies on the usage of the same combination of drugs against ovarian cancer. Of interest, we observed an increase in GSH in both PEO1 and PEO4 cells when treated with auranofin, suggesting that the cells may compensate the oxidative environment caused by the blockage of TrxR. However, when we combined auranofin with L-BSO, the levels of GSH were reduced, an effect that was associated with further lethality in the groups treated with auranofin-L-BSO versus the groups receiving auranofin alone. Together with the inhibition of GSH, combining AF with L-BSO led to more elevated levels of ROS than those caused by AF alone. Of particular interest, L-BSO alone was not able to increase ROS in PEO1 cells, but it did so in PEO4 cells. This difference in behavior of the cells in terms of L-BSO-mediated ROS production could be because PEO1 cells have a higher basal expression of the antioxidant TrxR than PEO4 cells. Regardless of the differences in behavior as monotherapy, when combined with AF, L-BSO increased the levels of ROS beyond those triggered by AF alone in both cell lines. This suggests that the combination of auranofin and L-BSO can be used as a chronic treatment against HGSOC to overcome platinum resistance by increasing the production of ROS which cannot be counteracted by any main active antioxidant system as both TrxR and GSH are inactivated.

## Conclusions

In summary, we report here that the gold complex auranofin is efficient to impair the functionality of HGSOC cells that are clinically sensitive or resistant to cisplatin (Figure 10). We further show that the drug was more efficient in killing HGSOC sensitive to cisplatin than the ones resistant to the platinum agent. The mechanism of cell death induced by auranofin involves inhibition of TrxR, depolarization of the mitochondrial membrane, production of ROS and caspase-associated apoptosis linked to DNA damage. This toxicity can be blocked by the antioxidant NAC indicating the relevance of ROS in the toxicity of auranofin. Furthermore, we provide evidence that in compensation for the pro-oxidant effect of auranofin, there is upregulation of GSH, which if blocked with L-BSO diminishes the antioxidant systems and potentiate the toxicity of auranofin in both platinum sensitive and resistant HGSOC cells. We anticipate that auranofin can be repurposed to chronically treat HGSOC after initial cytotoxic chemotherapy as a maintenance therapy alone or in combination with the antioxidant L-BSO.

**Figure 10.**
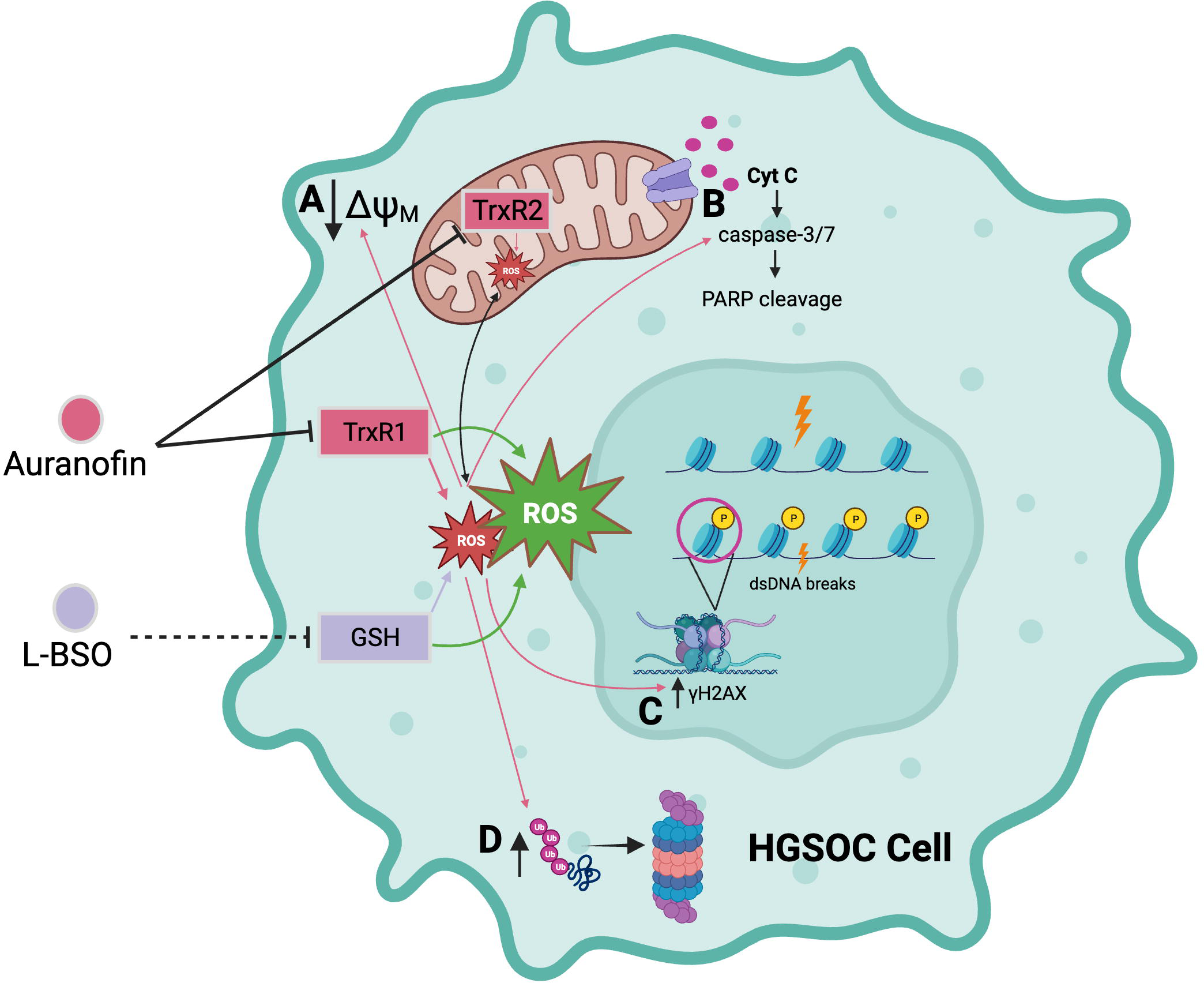
Auranofin (AF) inhibits the activity of the antioxidant enzymes thioredoxin reductase 1 (TrxR1) and 2 (TrxR2), inducing an increase in reactive oxygen species (ROS) in the high-grade serous ovarian cancer (HGSOC) cell. Increased ROS that was induced by AF caused; **A** decreased membrane potential in the mitochondrial membrane, **B** increased activation of the executor caspase-3/7 and cleavage of poly-ADP ribose polymerase (PARP), **C** double-stranded DNA (dsDNA) breaks and phosphorylation of the serine 139 residue of the histone H2AX (yH2AX), and **D** accumulation of poly-ubiquitinated proteins. L-buthionine sulfoximine (L-BSO) indirectly inhibits the production of the antioxidative protein, glutathione (GSH), resulting in an increase in ROS. The combination of AF and L-BSO results in a further increase in the production of ROS than the amount of ROS produced by each drug separately. Arrows in pink correspond to the effects caused by ROS induced by AF alone. Arrows in green signify a greater induction of ROS by the combination of AF and L-BSO. Created with BioRender.com.

## Supplementary Materials

Figure S1: Induction of apoptosis by auranofin is not prevented by caspase inhibition; Figure S2: Auranofin incudes accumulation of poly-ubiquitinated proteins, an effect prevented by NAC; Figure S3: original membranes containing detailed information of the uncropped immunoblot images.

## Author Contributions

Conceptualization, C.T. and F.A.; methodology, F.A. and E.T.; experiments with cisplatin, B.F.; formal analysis, A.G, E.Z., F.A., C.T.; MTT assays, E.M-J.; original draft preparation, F.A.; writing—review and editing, C.T.; critical revision, E.Z.; project administration, A.G.; All authors have read and agreed to the published version of the manuscript.”

## Institutional Review Board Statement

“Not applicable.”

## Informed Consent Statement

“Not applicable.”

## Data Availability Statement

The detailed data generated in the manuscript are openly available to the scientific community upon request.

## Acknowledgements

We are indebted to Millenium Pharmaceuticals for providing bortezomib (Velcade®) for preclinical research purposes.

## Conflicts of Interest

“The authors declare no conflict of interest.”

**Abbreviations**
ΔΨM: Trans-membrane potential
7-AAD: 7-Aminoactinomycin D
AF: Auranofin
ALL: Acute lymphoblastic leukemia
ATP: Adenosine triphosphate
BCA: Bicinchoninic acid
BRCA2: Breast cancer type 2 susceptibility protein
BZ: Bortezomib
CI: Combination index
CLL: Chronic lymphocytic leukemia
CO_2_: Carbon dioxide
DEVD: DNA-binding peptide sequence; substrate for caspase-3
DHE: Dihydroethidium
DNA: Deoxyribonucleic acid
DTNB: 5,5’-dithio-bis (2-nitrobenzoic acid)
DTT: Dithiothreitol
FBS: Fetal bovine serum
FDA: US Food and Drug Administration
GLOBOCAN: Global Cancer Observatory
GSH: Glutathione
H2AX: Histone variant H2AX
HCl: Hydrochloric acid
HEPES: 4-(2-hydrozyethyl)-1-piperazineethanesulfonic acid
HGSOC: High-grade serous ovarian cancer
HIV: Human Immunodeficiency virus
HRP: Horseradish peroxidase
IC50: Half-maximal inhibitory concentration
IgG: Immunoglobulin G
L-BSO: L-buthionine sulfoximine
MTT: 3-(4,5-dimethyliazol-2-yl)-2,5-diphenyltetrazolium bromide
NAC: N-acetyl-L-cysteine
NADPH: Nicotinamide adenine dinucleotide phosphate
NRF2: Nuclear factor erythroid 2-related factor 2
NSCLC: Non-small cell lung cancer
OD: Optical density
PARP: Poly-adenosine diphosphate (ADP) ribose polymerase
PBS: Phosphate-buffered saline
PE: Phycoerythrin
PeCy5: Phycoerythrin-Cyanine®5
PFA: Paraformaldehyde
PMSF: Phenylmethylsulfonyl fluoride
PS: Phosphatidylserine
ROS: Reactive oxygen species
SDS: Sodium dodecyl sulfate
TBS-T: Tris-buffered saline 0.1% Tween 20
TNB: 5’thio-2-nitrobenzoic acid
TrxR: Thioredoxin reductase
z-DEVD-fmk: Caspase-3 inhibitor

