## Supplementary figures and images for "Auranofin induces lethality driven by reactive oxygen species in high-grade serous ovarian cancer cells"

### Supplementary Figure 1

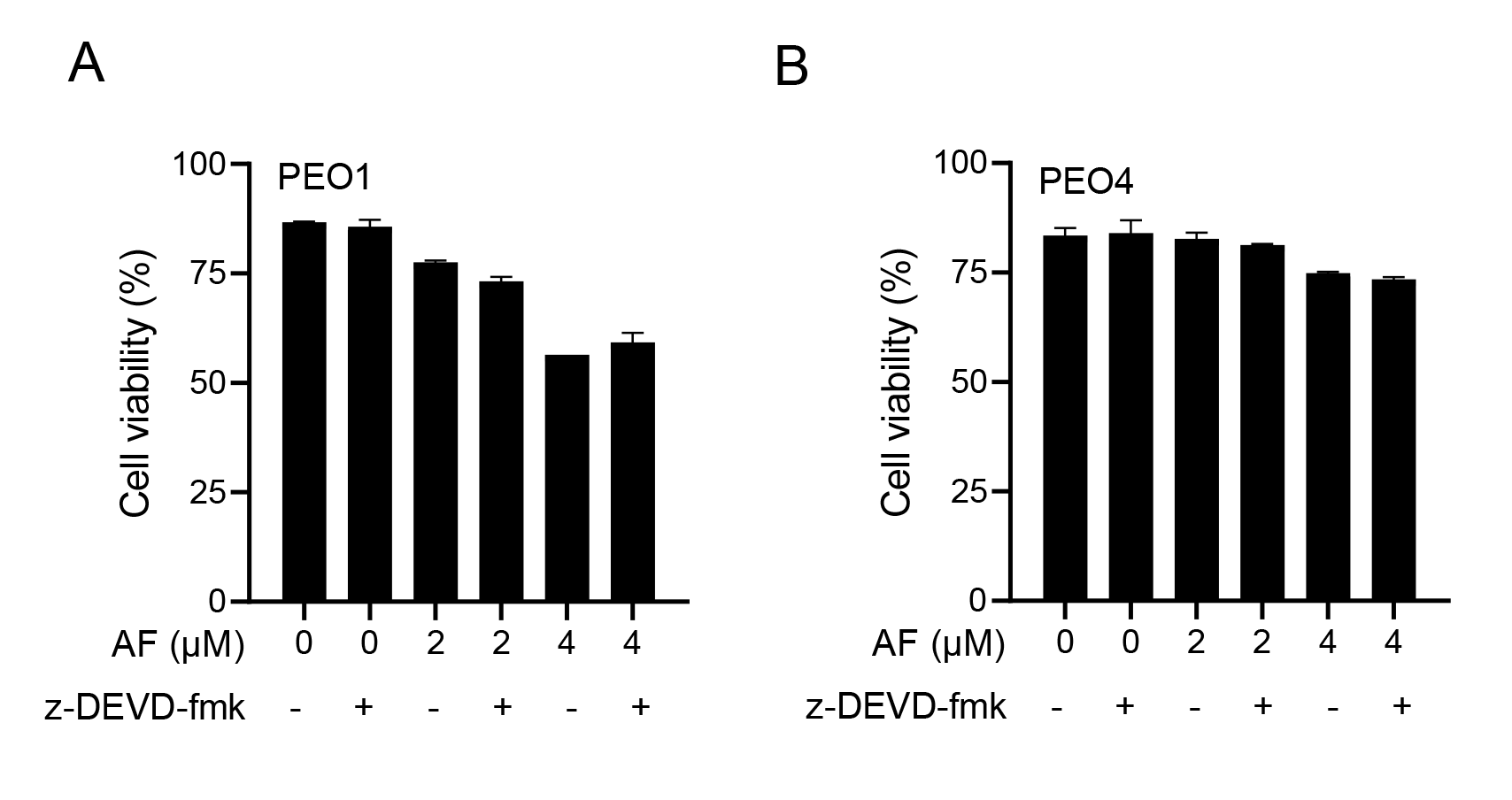

### Supplementary Figure 3

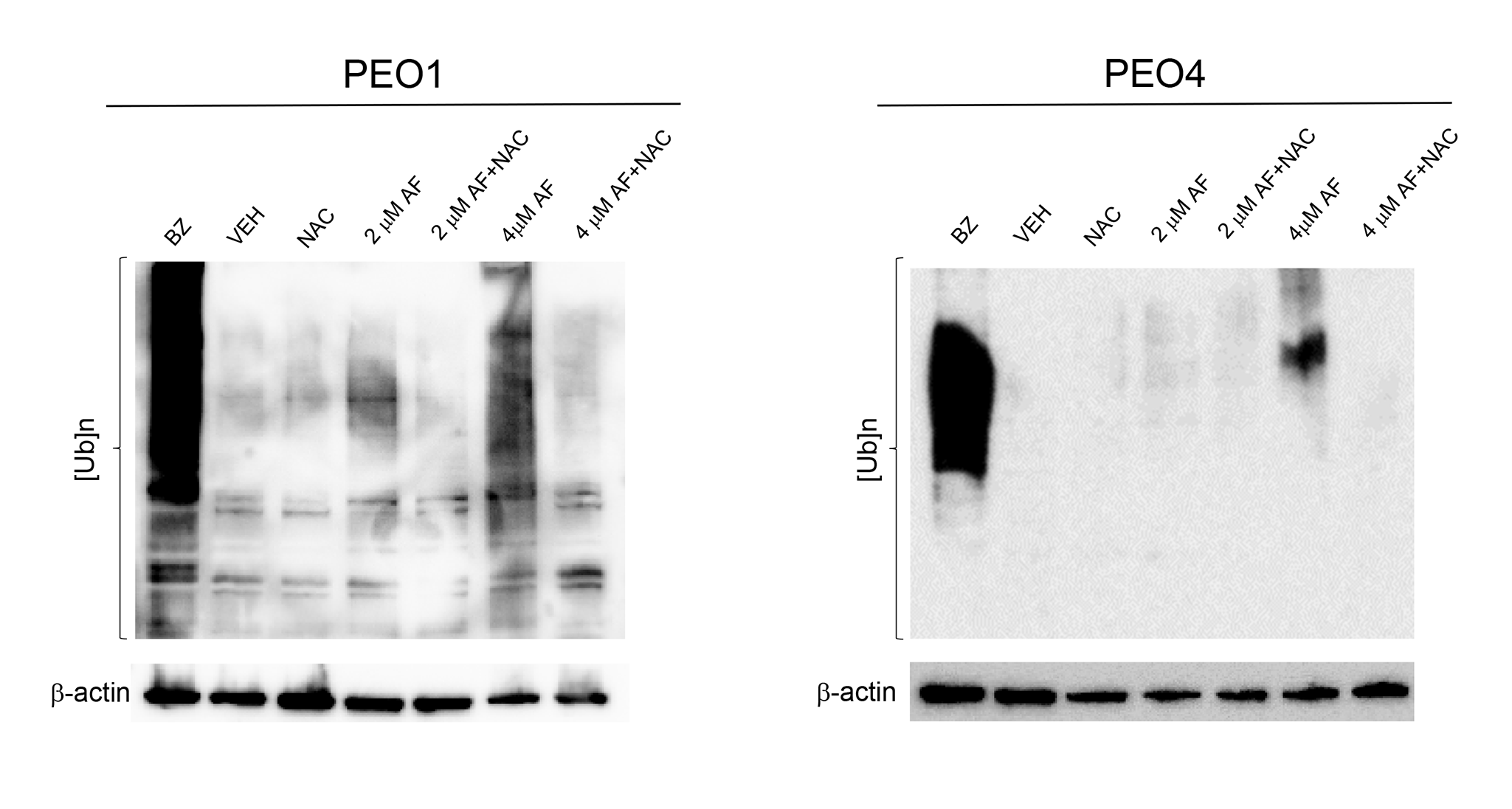
