## Supplementary Figure 3 for "Auranofin induces lethality driven by reactive oxygen species in high-grade serous ovarian cancer cells"

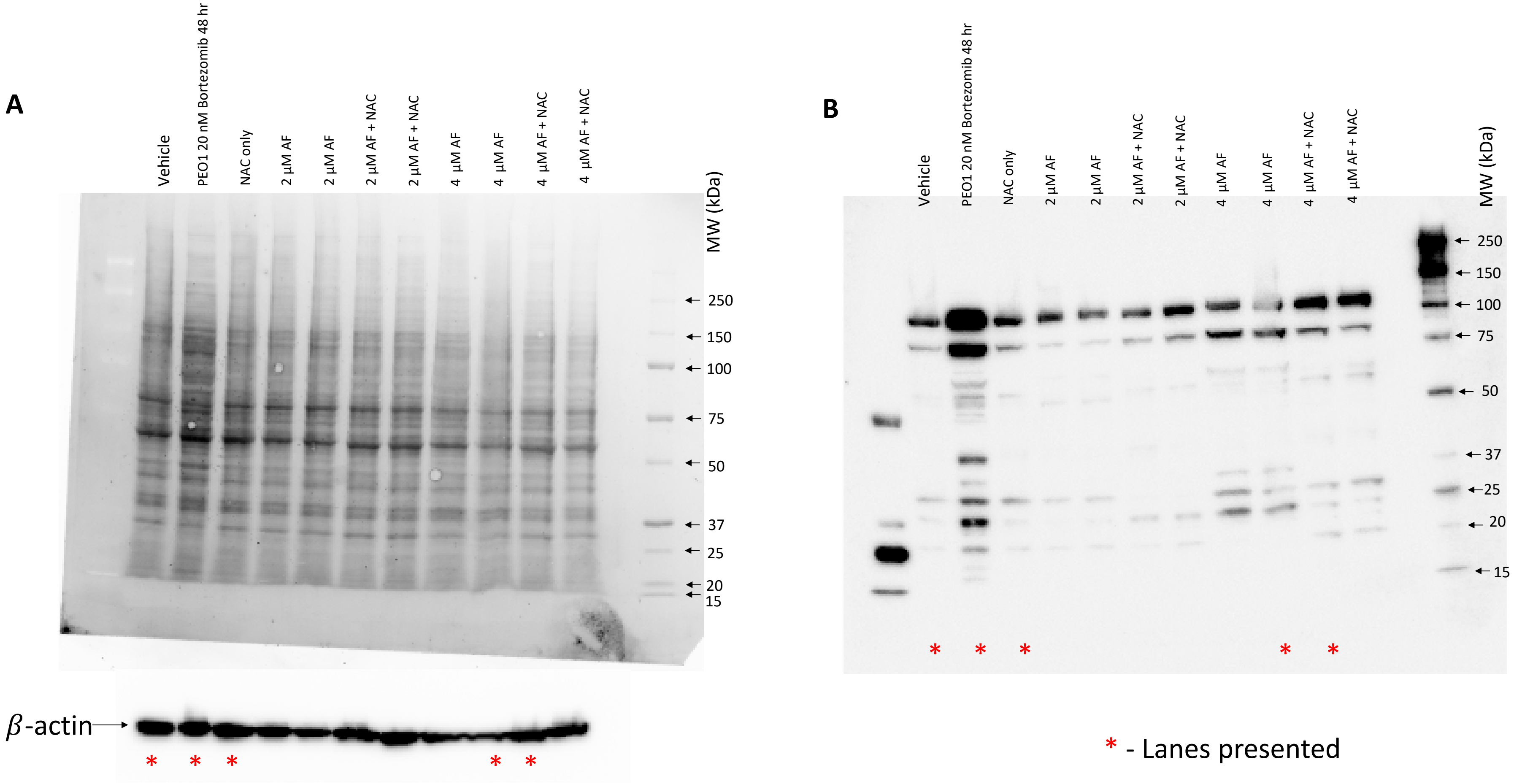


**Blot 1.** PEO1 cells were treated with the indicated concentrations of AF with or without the presence of 5 mM NAC for 24 hr. PEO1 cells treated with 20 nM bortezomib (Bz) for 48 hr were used as a positive control. Extracted proteins from the samples were run for 35 min at 200V on TGX stain free fast cast acrylamide gel (10 %). After the run, the gel was activated by UV light, and the proteins on the gel were transferred for 7 min using TransBlot Turbo. After the transfer, total protein load was visualized in the unstained membrane using BioRad ChemiDoc imager (A). After 1 h of blocking with 5% non-fat dry milk, the blot was incubated with anti-PARP antibody overnight at 4°C. (B). The blot was then incubated for β-actin (A). Data from this blot was presented in Figure 7B.

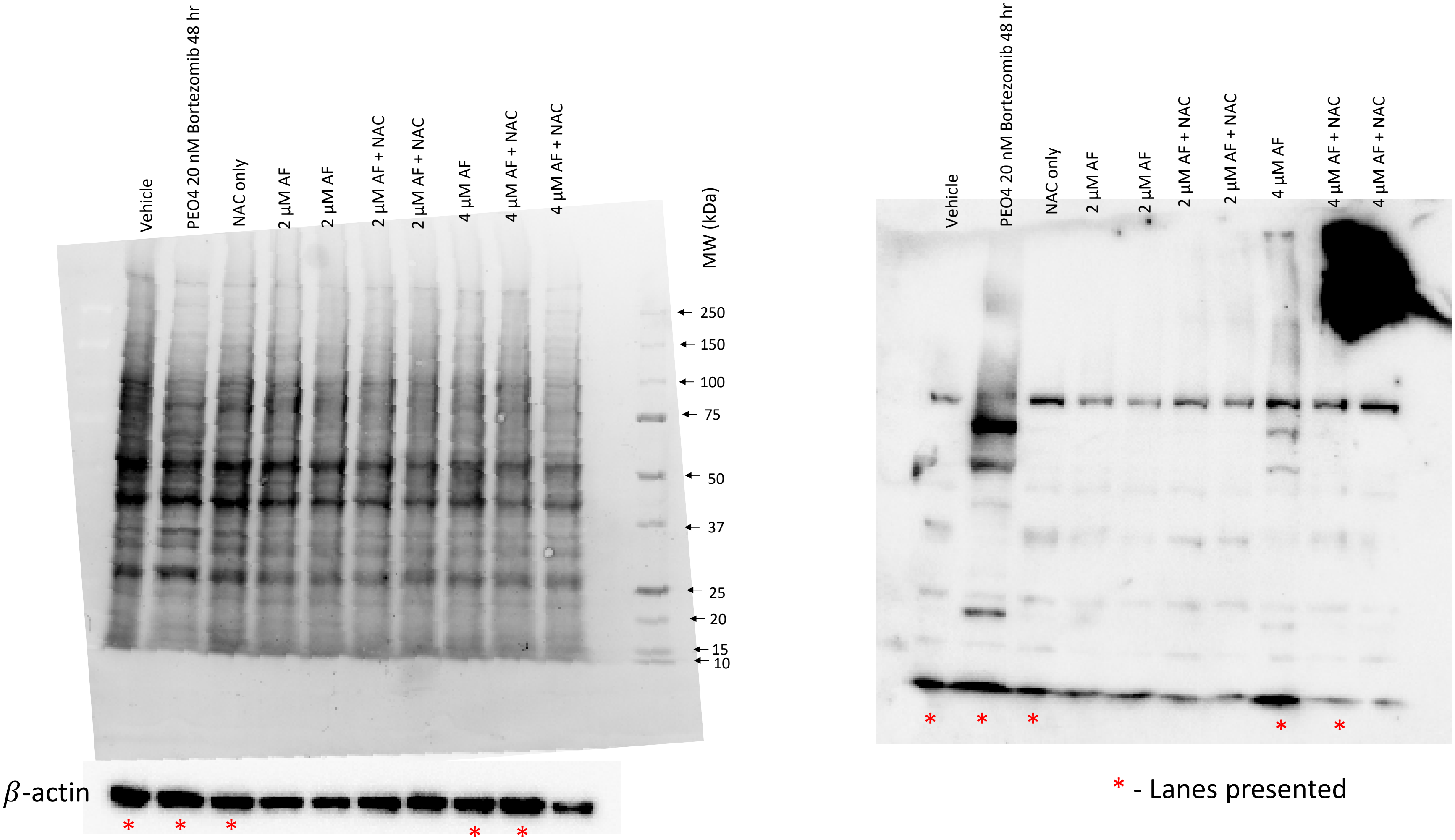


**Blot 2**. PEO4 cells were treated with the indicated concentrations of AF with or without the presence of 5 mM NAC for 24 hr. PEO4 cells treated with 20 nM bortezomib (Bz) for 48 hr were used as a positive control. Extracted proteins from the samples were run for 35 min at 200V on TGX stain free fast cast acrylamide gel (10 %). After the run, the gel was activated by UV light, and the proteins on the gel were transferred for 7 min using TransBlot Turbo. After the transfer, total protein load was visualized in the unstained membrane using BioRad ChemiDoc imager (A). After 1 h of blocking with 5% non-fat dry milk, the blot was incubated with anti-PARP antibody overnight at 4°C. (B). The blot was then incubated for β-actin (A). Data from this blot was presented in Figure 7D.

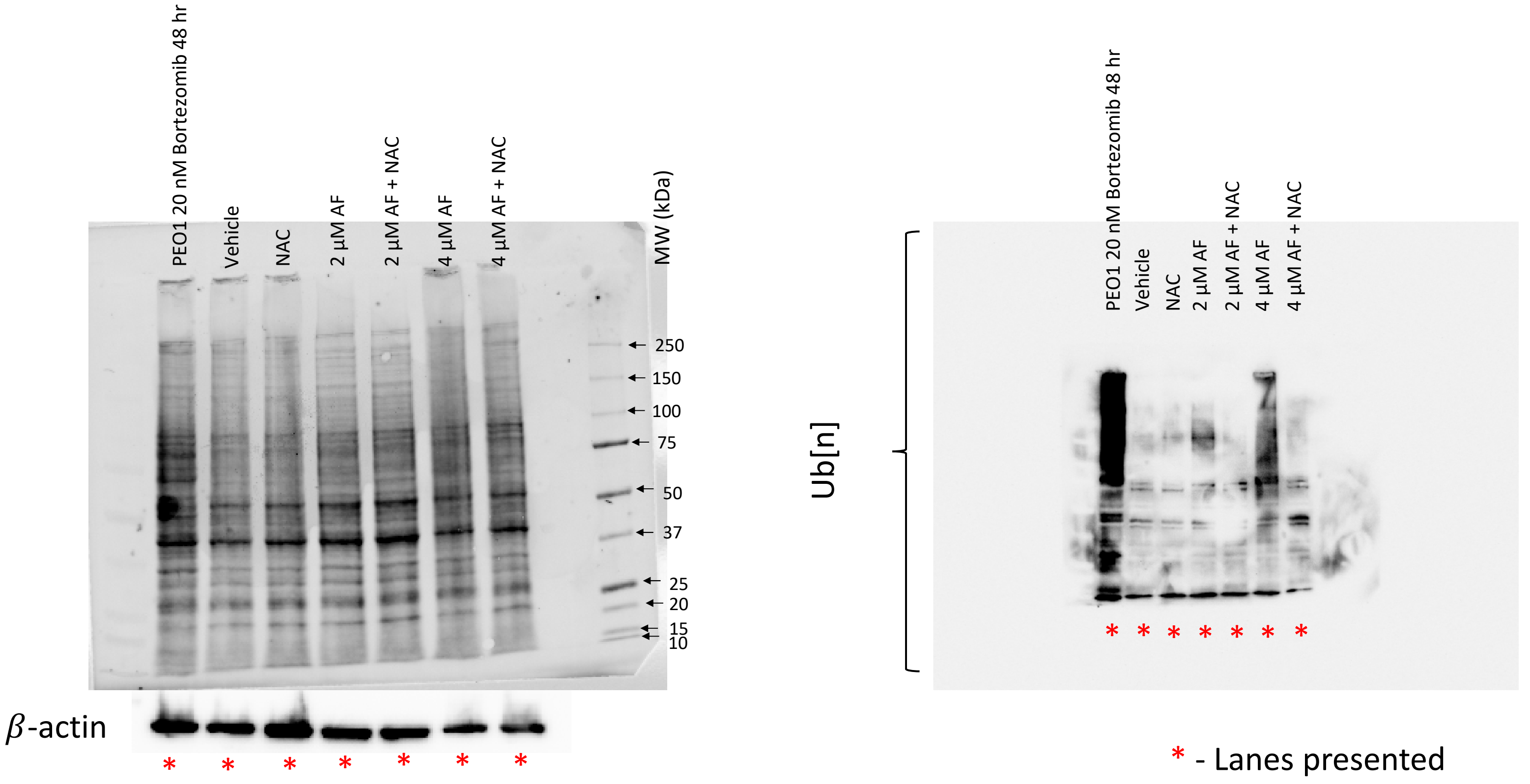


**Blot 3.** PEO1 cells were treated with the indicated concentrations of AF with or without the presence of 5 mM NAC for 24 hr. PEO1 cells treated with 20 nM bortezomib (Bz) for 48 hr were used as a positive control. Extracted proteins from the samples were run for 35 min at 200V on TGX stain free fast cast acrylamide gel (10 %). After the run, the gel was activated by UV light, and the proteins on the gel were transferred for 7 min using TransBlot Turbo. After the transfer, total protein load was visualized in the unstained membrane using BioRad ChemiDoc imager (A). After 1 h of blocking with 5% non-fat dry milk, the blot was incubated with anti-ubiquitin antibody overnight at 4°C. (B). The blot was then incubated for β-actin (A). Data from this blot was presented in Supplementary Figure 2.

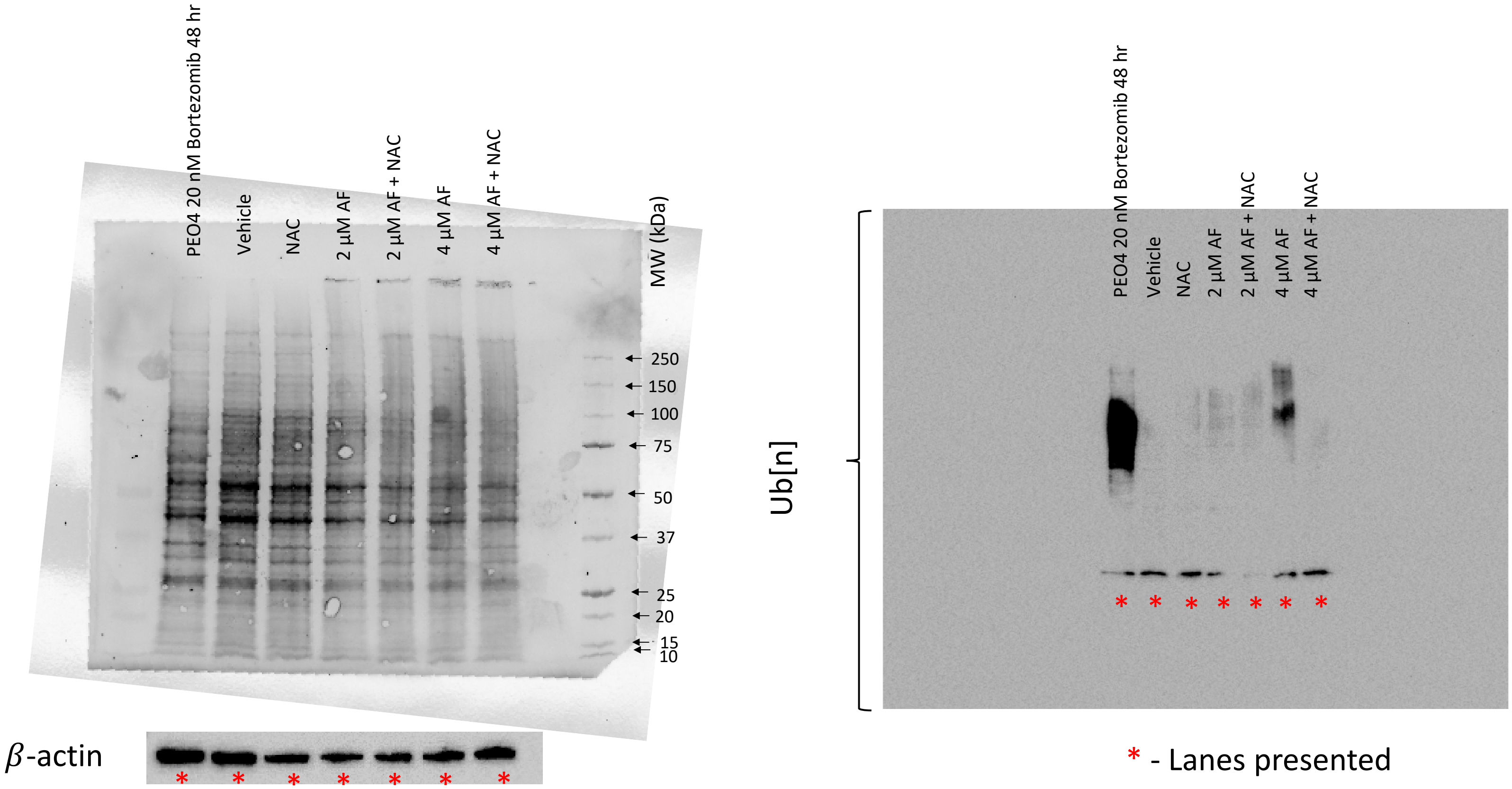


**Blot 4**. PEO4 cells were treated with the indicated concentrations of AF with or without the presence of 5 mM NAC for 24 hr. PEO4 cells treated with 20 nM bortezomib (Bz) for 48 hr were used as a positive control. Extracted proteins from the samples were run for 35 min at 200V on TGX stain free fast cast acrylamide gel (10%). After the run, the gel was activated by UV light, and the proteins on the gel were transferred for 7 min using TransBlot Turbo. After the transfer, total protein load was visualized in the unstained membrane using BioRad ChemiDoc imager (A). After 1 h of blocking with 5% non-fat dry milk, the blot was incubated with anti-ubiquitin antibody overnight at 4°C. (B). The blot was then incubated for β-actin (A). Data from this blot was presented in Supplementary Figure 2.
